# The cryo-EM structure of the chloroplast ClpP complex reveals an interaction with the co-chaperonin complex that inhibits ClpP proteolytic activity

**DOI:** 10.1101/2021.07.26.453741

**Authors:** Ning Wang, Yifan Wang, Qian Zhao, Xiang Zhang, Chao Peng, Wenjuan Zhang, Yanan Liu, Olivier Vallon, Michael Schroda, Yao Cong, Cuimin Liu

**Author notes:** these authors contributed equally to this work (Ning Wang, Yifan Wang).

## Abstract

Protein homeostasis in plastids is strategically regulated by the protein quality control system involving multiple chaperones and proteases, among them the Clp protease. We determined the structure of the chloroplast ClpP complex from *Chlamydomonas reinhardtii*by cryo-EM. ClpP contains two heptameric catalytic rings without any symmetry. The top ring contains one ClpR6, three ClpP4 and three ClpP5 subunits while the bottom ring is composed of three ClpP1_C_ subunits and one each of the ClpR1-4 subunits. ClpR3, ClpR4 and ClpT4 subunits connect the two rings and stabilize the complex. The chloroplast Cpn11/20/23 co-chaperonin, a co-factor of Cpn60, forms a cap on the top of ClpP by protruding mobile loops into hydrophobic clefts at the surface of the top ring. The co-chaperonin repressed ClpP proteolytic activity *in vitro*. By regulating Cpn60 chaperone and ClpP protease activity, the co-chaperonin may play a role in coordinating protein folding and degradation in the chloroplast.

## INTRODUCTION

Sophisticated pathways within cells control and regulate the biogenesis, trafficking and degradation of proteins to ensure protein homeostasis (proteostasis). These pathways belong to the protein quality control (PQC) system that involves the participation of various chaperones and proteases^1–5^. Molecular chaperones act as key components of the PQC system. They assist protein folding when new polypeptide chains emerge from ribosomes, when proteins have translocated through membranes, and when proteins become misfolded. Proteases are another important component of the PQC system that recognize and degrade substrate proteins which cannot fold or refold to the native state.

The molecular chaperone chaperonin is a tetradecameric cylinder formed by two rings stacked back to back, whose central cavity is used to shield a bound unfolded protein from the crowded environment and to assist its folding to the native state in an ATP-dependent process ^6^. A co-chaperonin complex seals the central cavity of group I chaperonins to form the isolated folding environment. In bacteria, this lid is formed by the homo-heptameric GroES co-chaperonins. In plastids, two types of co-chaperonin subunits exist, Cpn10 and Cpn20, the latter formed by two co-chaperonin domains in tandem. In addition to its role as a co-chaperonin for Cpn60, Cpn20 has been shown in *Arabidopsis* to mediate superoxide dismutase (FeSOD) activation independent of the Cpn60 chaperonin ^7^. In addition, the expression level of the *Cpn20* gene influenced the physiological function of ABA signaling during seed germination and promoted stomatal closure without Cpn60 involvement ^8, 9^. These results indicate that the Cpn20 protein might exhibit biochemical functions in addition to that as a co-factor for Cpn60 in protein folding.

In bacterial cells or endosymbiotic organelles, proteins are degraded by many types of proteases including serine-, aspartate-, and threonine-proteases ^10, 11^. The Clp protease is a conserved serine protease that consists of the ClpP core complex and assistant chaperones ^12^. The first crystal structure of the *E. coli* ClpP core complex revealed a barrel composed of two stacked heptameric rings, with one catalytic triad in each subunit ^13^. The degradation of protein substrates by the Clp protease in *E. coli* is assisted by two hexameric AAA+ (ATPase Associated with multiple cellular Activities) chaperones, ClpX and ClpA, both of which can bind to the ClpP core complex via one of its surfaces ^14^. Fueled by ATP hydrolysis, substrate proteins are unfolded by ClpX or ClpA and threaded into the central cavity of the ClpP core complex ^14^, where the substrate is degraded. Recent studies have explored the functional significance of the symmetry mismatch between the AAA+ chaperone hexamer and the heptamer of the ClpP core complex in *E. coli* ^15–18^. In particular, it has been proposed to induce a rotational movement between the AAA+ ATPase and the ClpP core and to participate in substrate transport ^16^.

In mitochondria, the ClpP core consists of 14 identical subunits that are similar to their bacterial homologs ^19^. In contrast, the ClpP core complex in chloroplasts is made up of three types of subunits termed ClpP, ClpR, and ClpT. ClpR subunits are homologous to ClpP but cannot contribute to catalysis because they lack one or more of the active residues of the Ser-His-Asp catalytic triad. ClpT proteins are completely different proteins of around 20 kDa that share homology with the N-terminal domain of the AAA+ chaperones. ClpT subunits have been proposed to link the two heptameric rings and thus maintain the stability of the tetradecameric ClpP core ^20–23^. In the green alga *Chlamydomonas reinhardtii*, the subunits of the chloroplast ClpP core complex are encoded by three *ClpP* genes (*clpP1*, *CLPP4*, *CLPP5,* the former chloroplast-encoded), five *ClpR* genes (*CLPR1-4*, *CLPR6*) and two *ClpT* genes (*CLPT3, CLPT4*) ^24, 25^. The plastidial *clpP1* gene produces ClpP1_H_, which can be further processed by unknown peptidases to generate ClpP1_N_, ClpP1_C_ and ClpP1_C’_ subunits which are part of the ClpP core complex ^26^. Translational attenuation of *clpP1* in *Chlamydomonas* led to the stabilization of misassembled photosynthetic enzymes ^27^, while its conditional repression caused serious autophagy responses and activated the protein quality control system ^28^. In *Arabidopsis* chloroplasts, the ClpP complex consists of five ClpP type subunits (ClpP3-6 and ClpP1), four ClpR type subunits (ClpR1-4) and two ClpT type subunits (ClpT1-2) ^20, 22, 29^. Loss of *ClpP5* gene function was embryo-lethal ^30^. Although the R-type subunits were considered unable to degrade substrate proteins, either *ClpR2* or *ClpR4* gene knockout resulted in delayed embryogenesis suggesting their functional importance ^30^. These results indicate that the Clp protease plays an essential role in maintaining protein homeostasis in chloroplasts, even though some subunits can be substituted by others. It was observed previously that the co-chaperonin Cpn20 interacts with the ClpP complex ^20^, but how this interaction takes place and whether it has functional implications are not known.

Recently, several structures of the Clp machinery composed of core complex and AAA+ chaperone have been resolved by cryo-EM. However these studies focused on bacterial Clp machineries, while there is no high-resolution structure of chloroplast Clp, yet ^15–17, 31^. In this study, we solved the cryo-EM structure of the *Chlamydomonas* chloroplast ClpP core complex. The complex is asymmetrical and polypeptide chains could be assigned to a known Clp gene. Furthermore, the co-chaperonin complex was observed to bind to the top ring of the ClpP core without inducing a major conformational change. We hypothesize that co-chaperonins act as regulatory factors in chloroplast proteostasis.

## RESULTS

### Co-chaperonins inhibit ClpP proteolytic activity via a direct interaction

In previous work, the ClpP complex has been purified from *Chlamydomonas reinhardtii* chloroplasts by Strep-tag affinity purification and the complex subunits have been assigned to P-, R- and T-type by mass spectrometry ^32^. Based on that work, we improved the purification strategy by combining affinity purification with multiple chromatography steps to obtain large amounts of highly purified *Chlamydomonas* ClpP complexes. The purified complexes were separated on a 12%-18% SDS gel and on a native gel and proteins were visualized by Coomassie staining (Figs. 1A and 1B). In the native gel, the purified ClpP complex migrated as a diffuse band with a molecular mass below the 820 kDa of oligomeric Cpn60 (Fig. 1B). By Asymmetric Flow Field Flow Fractionation (AFFFF) we calculated a molecular mass of 549 kDa for the ClpP complex, which is much larger than the 240 kDa of *E. coli* ClpP (Fig. S1A). Each visible protein band in the SDS gel was cut out and analyzed by mass spectrometry, and the bands identified as Clp or co-chaperonin subunits are marked (Fig. 1A) (Dataset 1). Though the migration pattern of Clp subunits was slightly different from the previous report due to difference in protein preparation and SDS-PAGE conditions, the same major Clp subunits were identified, except for ClpT3 (Table S1) ^32^. In addition to the Clp subunits, the three co-chaperonin proteins Cpn11, Cpn20, and Cpn23 were identified with notable abundance (Table S1)(Dataset 1). The presence of *clpP1* gene products ClpP1_H_, ClpP1_C_ and ClpP1_C’_ and of Cpn20 in the purified ClpP complex was confirmed by immunoblotting using antisera against the Strep-tag and against Cpn20 (Fig. 1C). It is of note that the two bands detected for both ClpP1_C_ and ClpP1_C’_ might result from the differentiated peptidase processing. To confirm the interaction of the Clp complex with the co-chaperonins, we performed a co-immunoprecipitation experiment using Cpn20 antiserum. As shown in Fig. 1D, Clp subunits ClpP1_C_, ClpP4, ClpR6, and ClpT4 were detected in the immunoprecipitate by immunoblotting. The interaction between the ClpP complex and chaperonins *in vitro* was further validated by size exclusion chromatography (Fig. 1E). Some of recombinantly produced Cpn20 co-migrated with the *Chlamydomonas* ClpP complex, and co-migration was even more marked with a mixture of the three co-chaperonins Cpn11/20/23. Note that Cpn20 was also detected after long time exposure in the ClpP-only fractions (CrClpP row) because the co-chaperonin complex co-purifies with endogenous ClpP. No co-migration of Cpn subunits was observed with the *E. coli* ClpP complex, and the *E. coli* co-chaperonin GroES co-migrated neither with *Chlamydomonas* nor *E.coli* ClpP. These results indicate that the interaction between the co-chaperonin and the ClpP complex can be reconstituted *in vitro* and is a specific feature of the chloroplast system. This conclusion is in line with the finding that *Arabidopsis* Cpn20 interacts with the *Arabidopsis* Clp complex *in vivo* ^20^.

**Figure 1.**
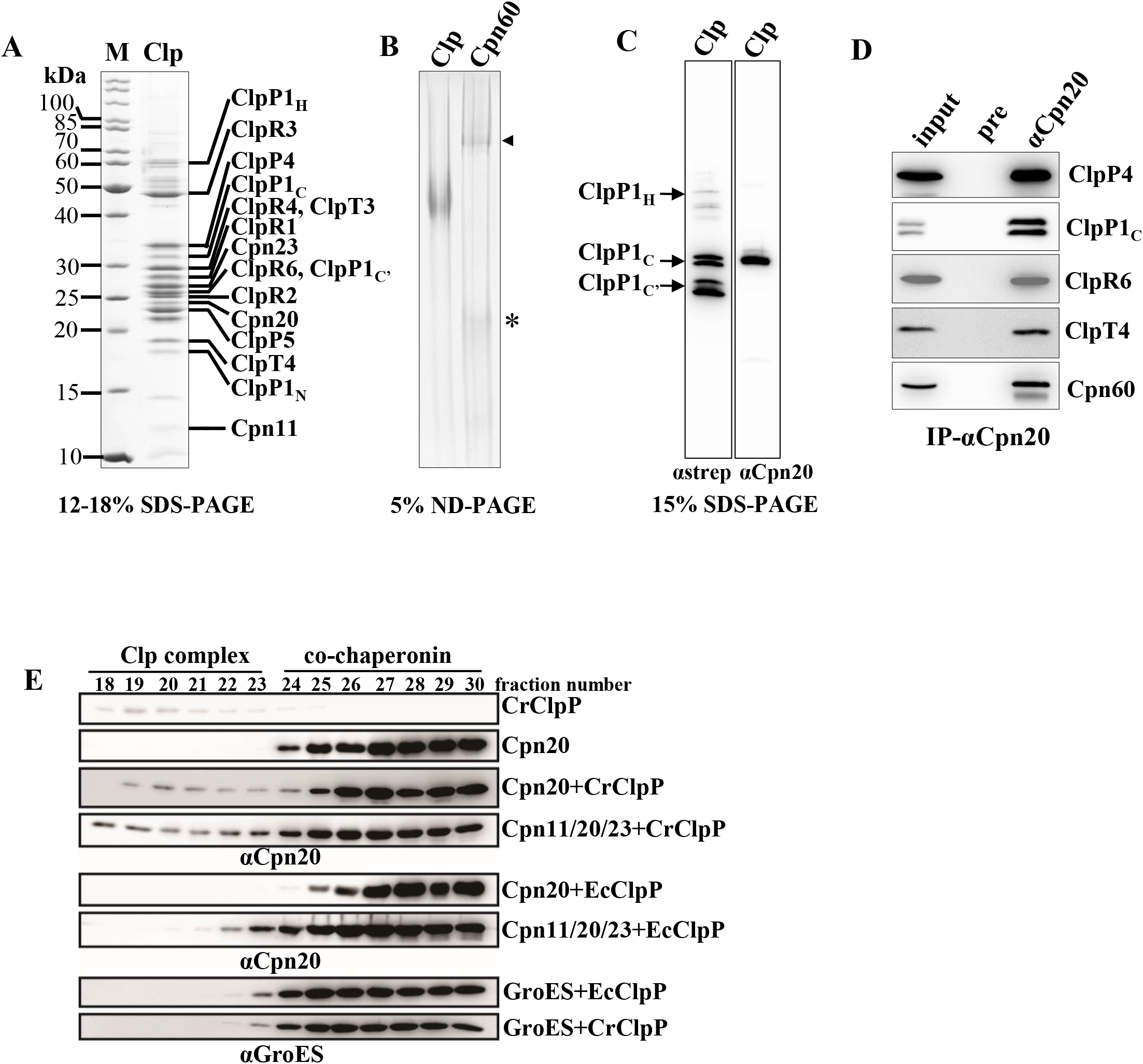
Co-chaperonins interact with CrClpP. **(A)** Purified *Chlamydomonas* ClpP complexes were separated on 12-18% SDS-polyacrylamide gels and visualized by Coomassie staining. Proteins in visible bands were identified by mass spectrometry and the bands identified as Clp or co-chaperonin subunits are marked. **(B)** Purified ClpP and Cpn60 complexes were analyzed on a 5% non-denaturing (ND) polyacrylamide gel and visualized with Coomassie staining. The arrowhead indicates the position of the oligomeric 820-kDa Cpn60 complex and the asterisk the position of the 60-kDa Cpn60 monomer. **(C)** Immunoblots analysis. Purified ClpP complexes were separated on a 15% SDS-polyacrylamide gel, transferred to nitrocellulose and immunodecorated with anti-strep and anti-Cpn20 antibodies. The Strep-antibody recognized the three gene products of the plastid-encoded *clpP* gene, ClpP1_H_, ClpP1_C_ and ClpP1_C’_. **(D)** Immunoprecipitation of Cpn20. Total cell lysates from the ClpP-Strep (#8) strain (input) were incubated with protein A-Sepharose beads coupled to antibodies of either preimmune serum (pre) or anti-Cpn20 serum. Precipitated proteins were analyzed by immunoblotting using antisera against ClpP4, the Strep-tag (detecting ClpP1_C_), ClpR6, ClpT4 and Cpn60. **(E)** Co-migration of ClpP core complexes and co-chaperonins in size exclusion hromatography. 1 μM ClpP protein complex was injected into a Superdex 200 PC 3.2/10 column after incubation with or without 2 μM co-chaperonin for 30 min at 4◻. Proteins were eluted with 20 mM MOPS-KOH, pH 7.5, 80 mM NaCl, 10 mM MgCl_2_, 10 mM KCl, 1 mM DTT, 10% glycerol. Then the corresponding fractions were analyzed by SDS-PAGE and immunoblotting using antisera against Cpn20 or GroES. The positions of the ClpP and co-chaperonin complexes are indicated at the top of the panel.

Beta-casein is a commonly used model substrate for the Clp protease (Caseino-Lytic Peptidase) ^33, 34^. To analyze protease activities of purified ClpP complexes, we performed protein degradation assays. ClpP of *E.coli* (EcClpP) could not degrade β-casein, unless 4 μM of the ClpP activator acyldepsipeptide (ADEP) was present in the reaction (Figs. 2A, 2B and S2A). ADEP increases the interaction between ClpP monomers, competes with the Clp ATPases for their binding sites on ClpP, and triggers a closed- to open-gate transition of the substrate entrance pore, which is otherwise tightly closed ^35, 36^. In contrast, purified *Chlamydomonas* ClpP was able to degrade β-casein without need for ADEP. ADEP slightly accelerated protein degradation by CrClpP, but only at a high concentration of 18 μM, not of 4 or 8 μM (Figs. 2A, 2B and S2A). Given the specific interaction between *Chlamydomonas* ClpP and the co-chaperonin complex (Fig. 1), we wondered whether co-chaperonins had an effect on ClpP proteolytic activity. Recombinant co-chaperonins Cpn20 and Cpn11/20/23 both slowed down the proteolytic activity of CrClpP (Figs. 2C and 2D), but co-chaperonin GroES had no effect (Figs. S2A and S2B). This inhibitory effect was overcome by the addition of 18 μM ADEP (Figs. 2C and 2D). CrClpP hydrolyzed β-casein into several fragments of lower molecular mass ranging from 15 to 20 kDa, which were not observed with EcClpP (Fig. S2A). Hence, casein degradation by CrClpP appeared less processive than by EcClpP+ADEP. Moreover, Cpn20 was still bound to the CrClpP complex after incubation with 18 μM ADEP (Fig. S2C), indicating that either ADEP and the co-chaperonin bind to different sites on the ClpP core complex or ADEP cannot expel the co-chaperonin from its binding sites. As tiny amounts of AAA+ chaperones might have co-purified with CrClpP, we cannot conclude that CrClpP is proteolytically active in the complete absence of AAA+ chaperones. However, treatment with hexokinase/glucose, which will remove all ATP from the buffer, had no effect on casein degradation (Fig. S2A), suggesting that CrClpP can degrade this substrate in a completely energy-independent process.

**Figure 2.**
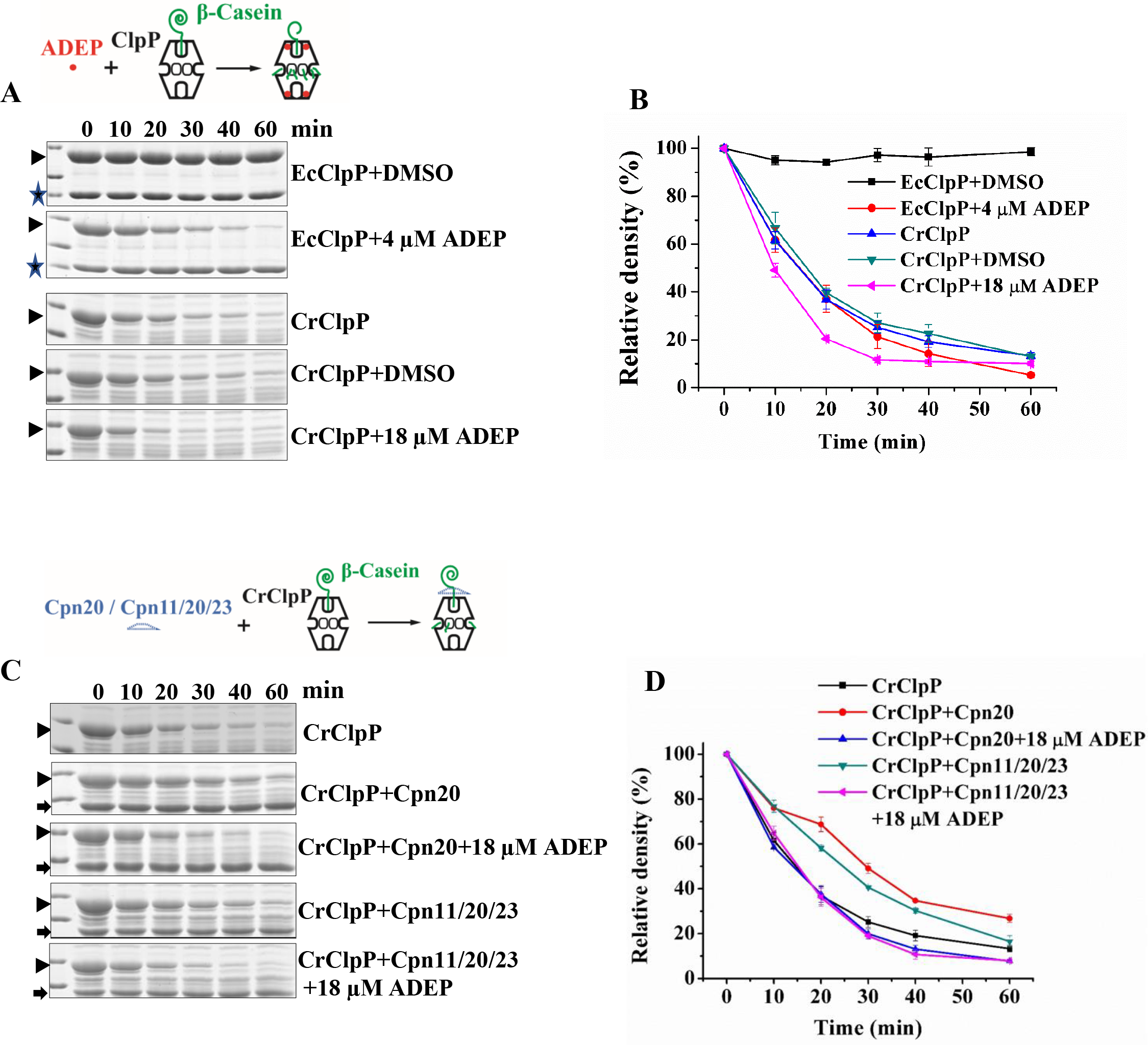
Co-chaperonins inhibit ClpP proteolytic activity. **(A)** and **(C)** Degradation of β-casein was monitored in reactions containing β-casein (16 μM), CrClpP (0.4 μM), EcClpP (0.4 μM), Cpn20 (0.4 μM), Cpn11/20/23 (0.4 μM) and ADEP dissolved in DMSO (4 or 18 μM) as indicated. Reactions were performed at 30C, aliquots were taken at the indicated time points, and analyzed via SDS-PAGE (15% gels) and Coomassie staining. Arrowheads indicate the position of β-casein, stars indicate the position of the EcClpP protein and arrows indicate the position of Cpn20. **(B)** and **(D)** Densitometric quantification of β-casein from the reactions shown in **(A)** and **(C)**, respectively. Shown are mean values from three independent replicates, error bars represent SD. Quantifications were made with Photoshop CS6.

### Cryo-EM structure of the purified ClpP complex and subunit assignment

To analyze structural features of the *Chlamydomonas* plastidic ClpP complex, we performed cryo-EM single particle analysis on the system. Purified ClpP complexes were applied to specimen grids and vitrified for cryo-EM analysis (Figs. S3 and S4). After 3D classification and iterative refinement, the optimally visualized particle groups were selected to generate two major groups, named ClpP-S1 and ClpP-S2, at resolutions of 3.3 Å and 3.6 Å, respectively. The main difference between these two groups was the appearance of a small cap in the latter particles (Figs. 3A-B and S3C, S4A, S4C-D). Focused 3D classification, particle re-extraction, re-centering and refinement of the cap, which was later identified as co-chaperonin Cpn11/20/23, generated particles with better structural features and more complete mobile loops at 4.8 Å resolution (Figs. S3C and S4B). We further combined the 4.8 Å Cpn11/20/23 and ClpP-S2 maps using the *vop maximum* command in Chimera and generated a ClpP-S2-composite map (ClpP-S2c)(Figs. 3B and S3C). The ClpP-S1 map displayed an obviously asymmetric structural conformation with a height of about 108 Å, a width of about 105 Å and a length of about 148 Å. The diameters of the central pore of ClpP-S1 are 28 Å and 20 Å for the top and bottom rings in the cut away views, respectively (Fig. 3A). This difference in pore size implied that there may exist a functional division between top and bottom ring in terms of substrate admittance. The cap in the ClpP-S2 particles increased the height to about 150 Å and the width to 118 Å, while the length was the same as ClpP-S1 (Figs. 3A and 3B). Overall, the resolution in the middle of the particles was higher than on the surface (Fig. S4D), especially for ClpP-S1 where the resolution in the particle core reached about 3 Å, which allows the identification of amino acid side chains.

**Figure 3.**
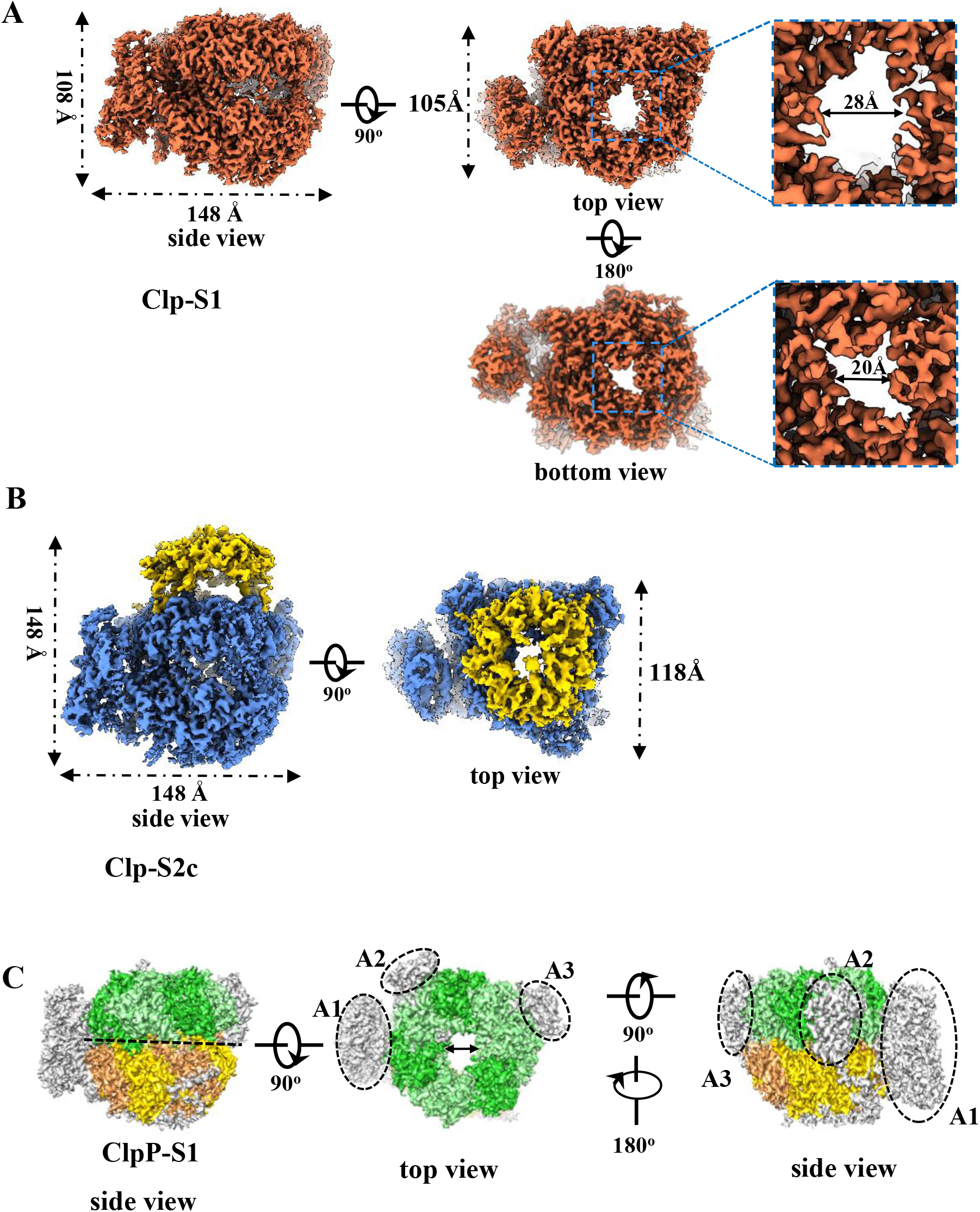
Cryo-EM structure of the *Chlamydomonas* ClpP complex. **(A)** Overview of ClpP-S1 particles. The dimensions are given next to the particles. The cut-away views of the central pore in the two rings of the ClpP core complex are shown. **(B)** Overview of ClpP-S2_C_ particles. The dimensions are given next to the particles. **(C)** Superimposition of EcClpP (PDB ID:1TYF) with density maps of CrClpP-S1. The subunits located in the top ring of EcClpP are shown in light or dark green and the subunits of the bottom ring are shown in yellow or orange. The dotted line indicates the interface between the two rings. Additional densities of CrClpP not overlapping with EcClp are shown in gray. The most prominent ones are labeled A1, A2 and A3.

To analyze the structural features of the CrClpP complex, the crystal structure of the *E.coli* ClpP complex (PDB: 1TYF) was manually docked into the CrClp-S1 density map (Fig. 3C) ^13^. The EcClpP structure fits well into the ClpP-S1 map around the central pore, with clear separation of double rings. Compared to the symmetrical EcClpP structure, three prominent additional densities appeared in ClpP-S1 that were labeled respectively as A1, A2 and A3. The A1 density is very large and locates adjacent to the interface of the two rings like a handle, while densities A2 and A3 appear to be more similar and locate to the peripheral region of the top ring.

In *Chlamydomonas*, three P-type (ClpP1, P4, P5), five R-type (ClpR1-4, R6) and two T-type (ClpT3, T4) subunits constitute the ClpP complex. ClpP1_H_ is processed by unknown peptidases to generate ClpP1_N_, ClpP1_C_ and ClpP1_C’_ subunits, which are part of the ClpP core complex. An alignment of amino acid sequences of P- and R-type subunits with EcClpP revealed a conserved region corresponding roughly to amino acids 27-175 in EcClpP (Figs. S5), which served to build the structures of the P- and R-type subunits (Fig. S6A). The structure of ClpP1_N_ could not be built after many trials. To assign the subunits into the ClpP core complex, the 14 subunits in the complex were designated as D1-D14 (Fig. 4A, left) and the individually built structures were manually fit into the central region of the ClpP-S1 map. Some specific sequences of ClpP and ClpR subunits were used for assigning them to the ClpP-S1 map (underlined in Fig. S5). Visualizations of map fitting of specific side chains allowed us to identify subunit locations in the Clp-S1 map (Fig. S6B).

**Figure 4.**
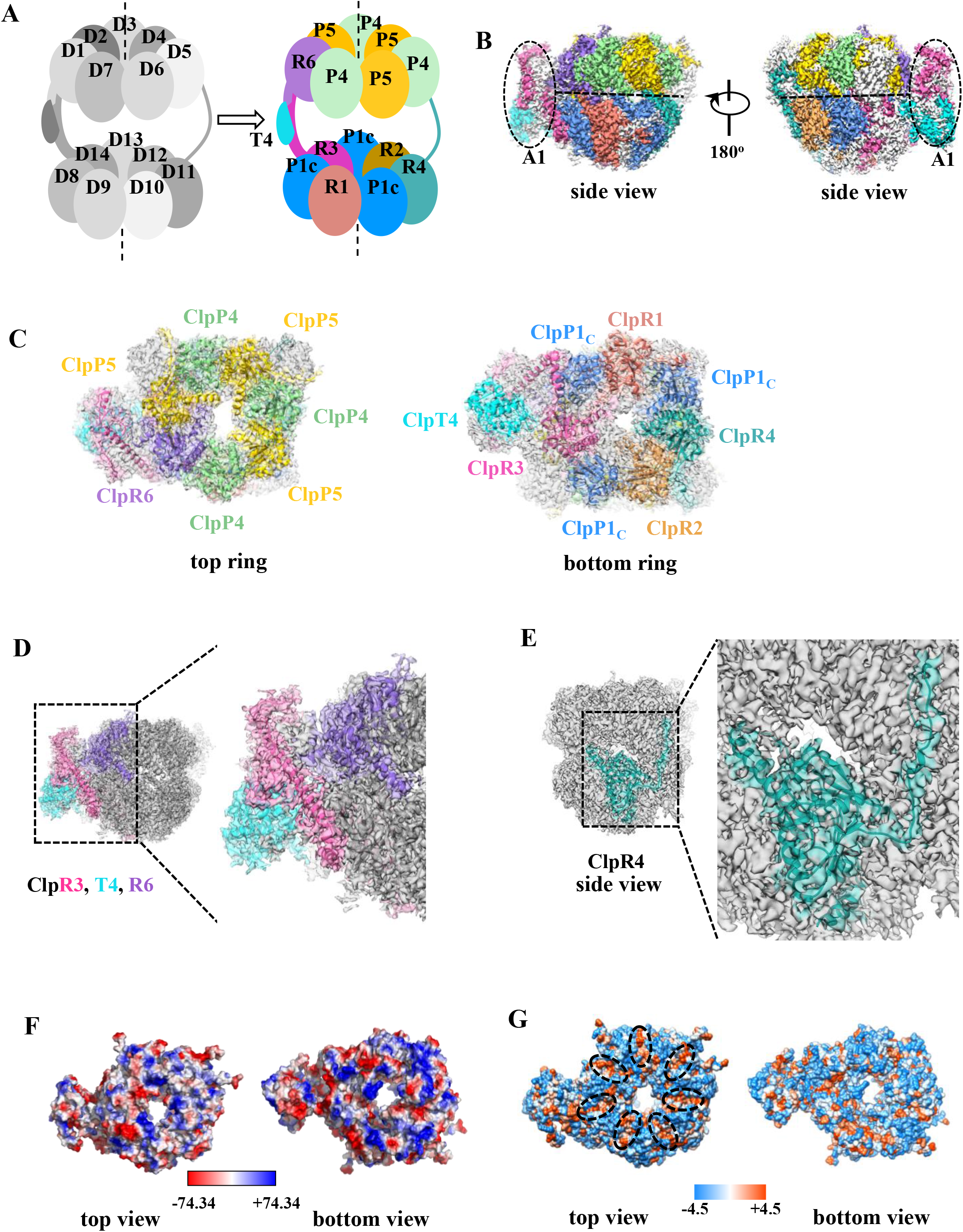
Properties of the *Chlamydomonas* ClpP core complex. **(A)** Cartoon presentation of the CrClp core complex. The subunits designated as D1-D14 are shown in the left grey cartoon and the corresponding assigned subunits are colored at the right. Subunit ClpP1_C_ occupies positions D8, D10, D13 positions. Subunit ClpP4 occupies positions D3, D5 and D7. Subunit ClpP5 occupies positions D2, D4 and D6. Subunits ClpR1, R2, R3, R4 and R6 occupy single positions at D9, D12, D14, D11 and D1 positions, respectively. **(B)** Density map of the ClpP core complex. Densities with assigned Clp subunits are shown in colors, densities that could not be assigned are shown in white. **(C)** Subunit composition and arrangement in the top and bottom rings of the ClpP core complex. **(D)** Assignment of Clp subunits to the additional density map A1. This region is omposed of full-length ClpT4 (cyan), the long C-terminus of ClpR3 (pink), and several amino acids from the C-terminus of ClpR6 (purple). The zoom-in view shows the fitting of these three subunits to the density. **(E)** Position of the ClpR4 subunit (turquoise) in the bottom ring of the ClpP core complex. The long C-terminus of ClpR4 protrudes to a region next to the top ring. The fitting of ClpR4 is shown in the zoom-in view. **(F)** Electrostatic potential of the ClpP core complex. Acidic amino acids are shown in red, basic amino acids in blue. **(G)** Hydrophobicity of amino acids at the ClpP core surface. Hydrophobic amino acids are shown in red, hydrophilic amino acids are shown in light blue. The seven hydrophobic binding clefts observed in the top ring of Clp-S1 are encircled by black broken lines.

After subunit assignment into the ClpP-S1 map, the individual subunit structures were built manually with continuous map densities. The structures of the N- and C-termini of some subunits could not be solved due to discontinuity in the densities. Information on the resolved sequences is summarized in Table S2. The superimposition of the individual subunit structure onto its corresponding electron density indicates that the models fit well into the corresponding maps (Fig. S7A). Overall, we could assign one ClpR6, three ClpP4, and three ClpP5 subunits to the top ring (designated as P-ring), and one of each ClpR1-4 and three ClpP1_C_ subunits to the bottom ring (Figs. 4A-C). Values of around 0.7 in the correlation coefficient (CC) chart corroborate our model-to-map fitting for each subunit (Fig. S7B). Still, some regions in the ClpP structure could not be solved, as shown by white regions in the core complex map density (Figs. 4B and S7C). Notice that no subunits or sequences could be fit into the additional A2 and A3 maps (Fig. S7C), including the long C-terminal sequences (Val196-Trp296) of the adjacent ClpP4 subunit, which remains unsolved. Conversely, no density could be ascribed to the large IS1 sequence characteristic of ClpP1_H_ ^24, 26^. The T4 subunit could be assigned to the A1 density, which is located adjacent to the interface of the two rings (Figs. 4B-C). The additional A1 map density, located at the interface of the two rings, is formed by a very long helix originating from the C-terminus of the ClpR3 subunit in the bottom ring, as well as by the T4 subunit and several amino acids from the ClpR6 subunit in the top ring (Fig. 4D). Some unassigned map density in A1 might be contributed by the C-terminus of ClpR3, and by ClpT3 or ClpP1_N_ subunits. The A1 region connects the two rings like a handle and can be assumed to stabilize the ClpP core complex. Moreover, the C-terminus of ClpR4 protrudes into a region next to the ClpP5 subunits in the top ring, and this might contribute to stabilize the core complex, as well (Fig. 4E).

The electrostatic potential is not equally distributed around the central pores of the two rings through which substrates enter the catalytic chamber, suggesting a differential affinity to substrates (Fig. 4F). The AAA+ chaperones interacts with EcClpP by protruding their flexible IGF loops into the hydrophobic clefts of ClpP which are formed at the interface of two subunits ^16^. Similarly, seven hydrophobic clefts were found to form at the surface of the top ring of ClpP by amino acids from two adjacent subunits. These hydrophobic clefts are arranged in a circular manner with seven-fold symmetry and were observed only on the top ring (Fig. 4G). However, we cannot completely rule out that similar hydrophobic clefts exist on the bottom ring, because the models of the three ClpP1c subunits in the bottom ring are not complete. A schematic model of the overall structure of the ClpP core complex with assigned subunits is shown in Fig. 4A.

### The cap on the top of the ClpP core complex is the co-chaperonin

We have shown that the co-chaperonin complex interacts with the ClpP core *in vivo* and *in vitro* (Fig. 1). Previous work from us and others showed that the authentic co-chaperonin complex in vivo consists of two Cpn20 subunits, one Cpn23 subunit, and one Cpn11 subunit ^37, 38^. Compared to Clp-S1, the ClpP-S2c map showed a dome-like density located on the top of the ClpP core complexes (Fig. 3, labeled A4 in Fig. S8A) which we attributed to the co-chaperonin. Since Cpn20 and Cpn23 each have two GroES-like domains, we split them into Cpn20-N, Cpn20-C, Cpn23-N, and Cpn23-C regions after cleavage of transit peptide ^39^ and performed a sequence alignment with Cpn11 and GroES (Fig. 5A). Sequences contributing to the roof of the dome-shaped co-chaperonin complex were present in GroES, Cpn23-N, Cpn23-C and Cpn20-C. This sequence was much shorter in Cpn11 and missing in Cpn20-N. Based on these specific roof characteristics we could assign Cpn11, Cpn23, and two Cpn20 subunits (Cpn20-1 and Cpn20-2) to the co-chaperonin map (Fig. 5B). The CC chart indicates a good model-to-map fitting for both Clp and co-chaperonin subunits (Fig. S8B) and the superimposition of the individual co-chaperonin subunit structure onto its corresponding electron density indicates that the models fit well into the corresponding maps (Fig. S8C). We further refined the model and fit five of the seven mobile loops into the densities that extended from the bottom of the co-chaperonin dome (belonging to the Cpn20 and Cpn11 subunits, Fig. 5B, left) (Fig. 5B, left). No densities were observed for the loops of the Cpn23 subunit, although the sequences are conserved (Fig. 5A).

**Figure 5.**
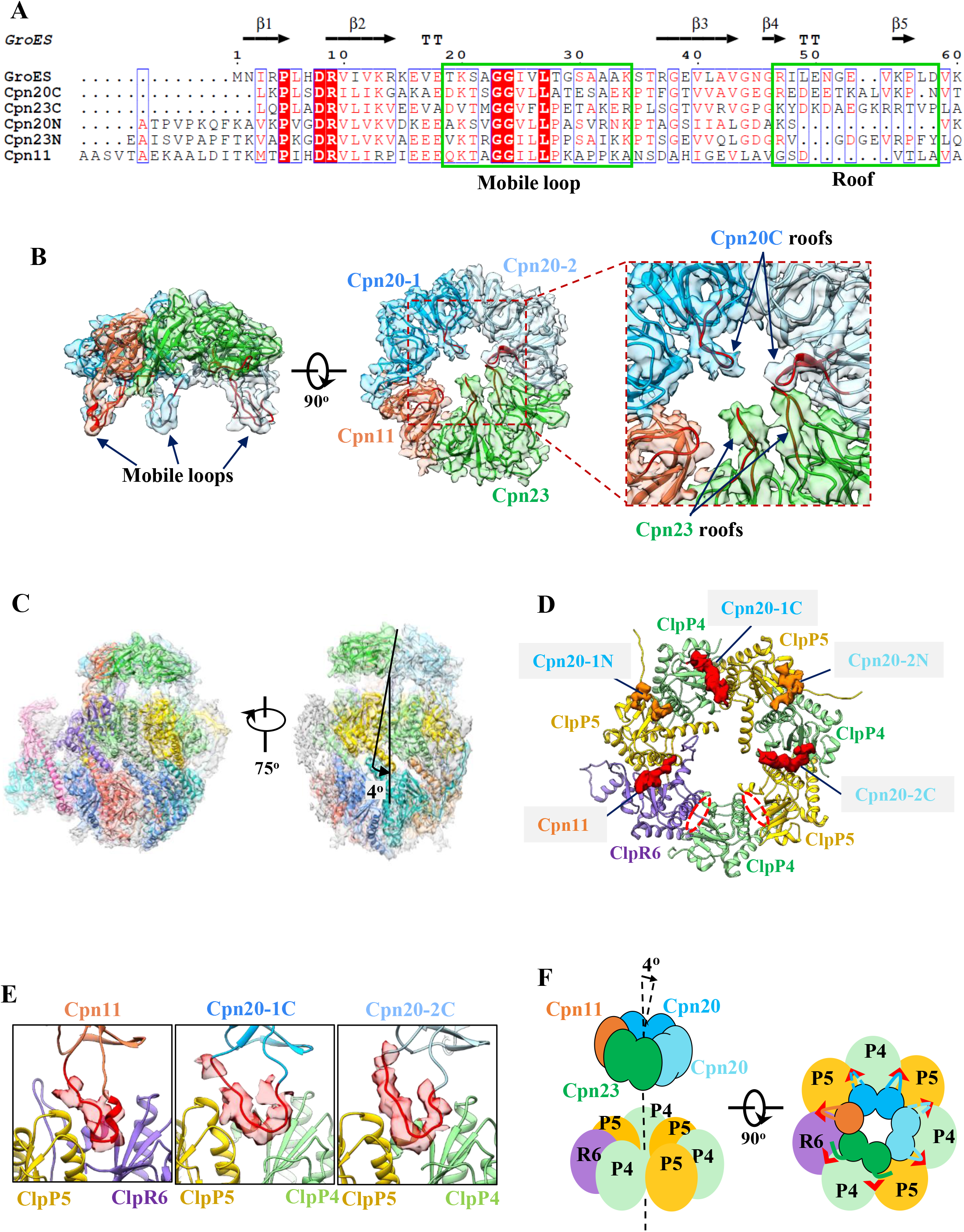
Cryo-EM structure of the *Chlamydomonas* ClpP-Cpn11/20/23 complex. (A) Sequence alignment of Cpn11, Cpn20-N, Cpn20-C, Cpn23-N, Cpn23-C and GroES. N and C indicate N- and C-terminal GroES-like domains of Cpn20 and Cpn23. The sequence regions of mobile loop and roof are indicated with green boxes. **(B)** Superimposition of the Cpn11/20/23 model and the cryo-EM density map. The mobile loops extending from the bottom of the co-chaperonin are shown (left). The enlarged picture shows the roof regions of Cpn23 and Cpn20C. **(C)** Interaction between ClpP and co-chaperonin. The central axes of co-chaperonin and CrClpP are given as black lines. The tilted angle between the axes of co-chaperonin and CrClpP complex has a value of 4°. **(D)** Top view oo the extra densities located in the hydrophobic clefts at the top ring of ClpP. The densities colored in red were confidently identified as mobile loops from Cpn11 and Cpn20C. The densities colored in orange might be the mobile loops from Cpn20N. The regions encircled with a broken red line are the two remaining clefts in which no additional densities could be identified. **(E)** Insertion of the mobile loops of Cpn11 and Cpn20C into the hydrophobic clefts at the surface of the top ring of ClpP. **(F)** Cartoon presentation of the interaction between the top ring of ClpP and Cpn11/20/23. Red arrowheads indicate the hydrophobic clefts. High-confidence mobile loops inserted into the clefts are indicated by solid lines, low-confidence mobile loops inserted into the clefts are indicated by broken lines.

Inspection of the map revealed that the co-chaperonin cap is tilted by about 4° relative to the ClpP symmetry axis (Figs. 5C and 5F). This tilt increases the distance between ClpP and Cpn23 versus ClpP and Cpn20, suggesting an intimate interaction at the Cpn20 side. A similar interaction, but with a tilt angle of 11°, has been observed for the asymmetrical EcClpP with its AAA+ chaperone ClpX ^16^. While that interaction is accompanied by a symmetry mismatch, this is not the case for the interaction between co-chaperonin and ClpP, which share a 7-fold symmetry. In *E.coli*, the IGF loops of ClpX insert into hydrophobic clefts located at the surface of EcClpP ^15^. Close inspection of the density map of the ClpP core complex in ClpP-S2c clearly revealed five extra densities in the hydrophobic clefts of the top ClpP ring (red and orange in Fig. 5D). Three of these densities were confidently identified as mobile loops stemming from Cpn11, contacting the ClpP5/R6 interface, and Cpn20-1C and Cpn20-2C, both contacting a ClpP5/P4 interface (red in Fig. 5E). The mobile loops from Cpn20-1N and Cpn20-2N might contact ClpP clefts as well, but at a slightly outward location, which could account for two more extra densities observed in clefts of the top ClpP ring (orange in Fig. 5D; Fig. S8D). An assignment of the densities in the two cleft regions was complicated by the close-by unassigned A2/A3 densities. For the two remaining hydrophobic clefts, in register with Cpn23, no extra map densities were observed (broken red lines in Fig. 5D), indicating that there is no direct interaction between Cpn23 and the ClpP core. In summary, the co-chaperonin appears to interact with the ClpP core by inserting at least three mobile loops into hydrophobic clefts on the surface of the top ring of ClpP (Fig. 5F), similar to the interaction of EcClpP with its AAA+ chaperone ClpX. Notice that the same loops of the co-chaperonin mediate its interaction with the Cpn60 chaperonin ^6^. The interactions of the co-chaperonin with ClpP, the co-chaperonin with Cpn60, and ClpP with AAA+ chaperones are thus mutually exclusive.

## DISCUSSION

### The chloroplast co-chaperonin has multiple functions

Many lines of evidence indicate that Cpn20 functions as a co-chaperonin for Cpn60 to assist protein folding in chloroplasts ^38, 40–42^. However, Cpn20 appears to exhibit functions independent of the Cpn60 chaperonin: in Arabidopsis, Cpn20 has been shown to play roles in abscisic acid (ABA) signaling and the activation of iron superoxide dismutase (FeSOD) ^7–9^. Here we show that a fraction of the *Chlamydomonas* chloroplast Cpn11/20/23 co-chaperonin complex co-purified with the ClpP core complex, and this interaction was robust enough to withstand several chromatography purification steps (Fig. 1). This interaction was shown before in *Arabidopsis* ^21^. The complexes between co-chaperonin and ClpP were confirmed by cryo-EM and *in vitro* reconstitution, and the co-chaperonin was shown to slow down the proteolytic activity of chloroplast ClpP (Figs. 2B and 5). The co-chaperonin caps the top-ring of the ClpP complex like a dome, similar to its interaction with Cpn60. Thus, the co-chaperonin might inhibit ClpP activity by blocking the entrance of unfolded proteins into the central cavity of the ClpP core complex via the pore of the top ring. The co-chaperonin complex interacts with ClpP and the Cpn60 chaperonin via the same mobile loops extending from the bottom of the dome (Fig. 5E). Therefore, the Cpn20/23/11 co-chaperonin complex appears to play a dual role in chloroplast protein quality control. Since its interaction with the Cpn60 chaperonin contributes to protein folding, the binding of the co-chaperonin to ClpP would simultaneously limit the efficiency of protein folding and inactivate a substantial part of the chloroplast’s degradation capacity. Releasing the co-chaperonin from ClpP would activate both activities at the same time, which might be important upon rapid challenges of proteostasis by environmental changes. In future studies, it will be interesting to quantify the relative stoichiometry of the chaperonin, co-chaperonin, and ClpP complexes in the *Chlamydomonas* chloroplast and their interactions under various growth or stress conditions. It is of note that the very abundant RbcL subunit of Rubisco is a substrate for both chaperonin and ClpP, and that translational attenuation of *clpP1* is lethal in the presence of *rbcL* mutations preventing its folding ^27^. While we initially postulated that the lack of sufficient ClpP would cause poisoning of the chaperonin by an unfoldable substrate, we must now also consider an additional hypothesis, that the mutant RbcL, by increasing the residence time of the co-chaperonin on and ever-busy Cpn60, would deregulate ClpP proteolysis.

The fact that the three Cpn60 subunits were also detected by mass spectrometry in our preparation (Dataset 1) raises another possibility, namely that ClpP and Cpn60 interact to regulate chloroplast protein homeostasis. Non-foldable substrate proteins, released from the chaperonin, need to be recognized by Clp to be degraded to prevent poisoning of the chaperonin. The co-chaperonin, by maintaining its interaction with the non-foldable substrate, could help direct it to the protease. In line with this idea, a cooperation between Trigger Factor, an ATP-independent ribosome-associated chaperone, and ClpXP was recently reported to promote the degradation of some substrates ^43^.

### Molecular architecture of chloroplast ClpP

The first structure of the *E. coli* ClpP core complex, obtained by X-ray crystallography, revealed 14 identical subunits in two stacked rings ^13^. The ClpP structures solved afterwards all consisted of a single or two types of ClpP subunits. In contrast, the chloroplast ClpP core complex solved here combines 10 different subunits, either of the P-type (harboring a functional catalytic site) or R-type (lacking catalytic residues) or T-type (homologous to the N-domain of AAA+ chaperones), distributed with uneven stoichiometry between two rings of different subunit composition. The subunit composition of the two rings of plastid ClpP from *Arabidopsis* was reported previously by the van Wijk and Clarke groups ^20, 22, 30, 44, 45^. The designated P-ring contains ClpP3, 4, 5 and 6 in a 1:2:3:1 ratio, while the designated R-ring consists of ClpP1 and ClpR1, 2, 3 and 4 in a 3:1:1:1:1 ratio. It was suggested that the two rings might exhibit different proteolytic capacities because only three catalytic subunits are present in the R-ring compared to seven in the P-ring. In the *Chlamydomonas* ClpP core complex reported here, the top ring consists of three ClpP4, three ClpP5 and one ClpR6 subunits. Because algal ClpP4 forms a sister clade to land plant ClpP3/ClpP4, and ClpR6 is clearly derived from ClpP6 by loss of a catalytic residue ^32^, the top ring of *Chlamydomonas* is clearly homologous to the *Arabidopsis* P-ring. Similarly, the bottom ring, with three ClpP1c and one each of ClpR1, 2, 3, and 4 subunits (Fig. 4C) can be called the R-ring. Because the diameter of the protein entrance pore in the P-ring is much larger than that in the R-ring (Fig. 3A), we propose that the former is the functional entry site of substrates. Only the P-ring was observed to interact with the co-chaperonin (Fig. 5), in line with a central regulatory role for this interaction. Our structure supports the notion that substrates are degraded inside a unique central cavity by a variety of active sites, possibly showing distinct chemical specificities as in the proteasome ^46^. Each subunit within each ring may also be functionally unique in terms of substrate selection, delivery and unfolding, and of interaction with the co-chaperonin. A good example is the specific interactions of the ClpP4/5 clefts with Cpn20 subunits (Fig. 5E, S8D).

Interesting differences can be noted between algal and land plant ClpP. With only 6 active subunits in its P-ring, the Chlamydomonas enzyme is slightly less asymmetrical than that of land plants in terms of catalysis (both have 3 catalytic sites in the R-ring). Importantly, the inactive ClpR6 subunit of the P-ring contributes, together with two other alga-specific features, namely the C-terminal extension of the R-ring ClpR3 and ClpT4, to the A1 side density that connects the two rings. In land plants, T-type Clp subunits were suggested to stabilize the ClpP core by interacting with both rings ^20, 21^, and it will be interesting to see if they form a structure similar to the A1 side-mass. Note however that the *Chlamydomonas* ClpT3/4 are vastly different from the ClpT1/2 of land plants ^32^, to the extent that they may not even be orthologous. Given that the N-terminal domain of AAA+ chaperones, to which ClpTs are homologous, play a role in substrate selection, the possibility arises that their dual structural/functional role was acquired several times in the evolution of the plastid Clp system. Furthermore, the C-terminus of ClpR4 protrudes to the P-ring, which also might stabilize the ClpP core complex. The massive contacts between the two rings may explain the high stability of the ClpP core, exemplified by our observation that the two-rings could not be separated by high salt treatment, in contrast to its homologs in *E. coli* and *Arabidopsis* plastids ^45, 47^.

### Proteolytic activity of the ClpP core might be regulated by the tilted interaction with the co-chaperonin complex

Structures of several different AAA+ chaperones with their ClpP cores have emerged recently, providing detailed information on the six-seven symmetry mismatch between the two complexes, and how it impacts their interaction dynamics ^15, 16, 18^. ClpX is tilted by 11° without major conformational changes upon binding to ClpP. As a result, the symmetry axes of the protease and the AAA+ chaperone are not aligned so that the translocation pathway for unfolded peptides is not straight but twisted. We also found a tilt of 4° between the axes of the co-chaperonin and the ClpP core (Fig. 5C). However, the fact that a single type of particle was obtained, with specific interactions between co-chaperonin and Clp subunits, together with the absence of a symmetry mismatch, suggest that the two complexes do not rotate and that the co-chaperonin instead operates like a cap stably sealing the entrance to the ClpP core.

Because none of the three known chloroplast AAA+ chaperones (ClpC1, ClpD1, ClpB3) were co-purified with ClpP in this and previous studies, they might not cooperate with ClpP in the chloroplast as they do in bacteria and mitochondria. Since the purified ClpP core complex was able to degrade the model substrate casein *in vitro* with no need for addition of ADEP, it appears possible that chloroplast ClpP can degrade proteins *in vivo* without the assistance of any type of chaperone. Regulation of protease activity may be carried out by the co-chaperonin or by small-molecule activators acting like ADEP.

## EXPERIMENTAL PROCEDURES

### Construction of expression plasmids for EcClpP and cochaperonins

The *ClpP* gene sequence from *E. coli* was amplified by PCR on genomic DNA and cloned into the pHUE vector with restriction enzymes to generate *EcClp*-pHUE. The produced EcClp protein contains ubiquitin and a 6×His tag at its N-terminus that facilitated subsequent protein affinity purification. The forward and reverse primers were 5’-GCGGATCCATGTCATACAGCAGCGGCGAACG-3’ and 5’-CCCAAGCTTTCAATTACGATGGGTCAGAATCGAATCGACCAG-3’ containing BamHI and HindIII restriction sites, respectively. The construction of the co-chaperonin expression plasmids, GroES-pET11a, CrCPN20-pQlinkT and CrCPN11/ 20/23-pQlinkT, were described earlier ^37^.

### *Chlamydomonas reinhardtii* strains and growth conditions

Chlamydomonas strain ClpP1-strep (#8), used for the purification of the Clp complex via Strep-tag affinity purification, has been describe previously ^26^ and is freely available upon request. The strain was kept on solid TAP medium containing 100 μg/ml of spectinomycin at 25◻ under continuous illumination (40 μmol photo m^−2^ s^−1^) or a photoperiod rhythm (12 h light/12 h dark).

### Purification of the ClpP complex from *Chlamydomonas reinhardtii*

Purification of the ClpP complex from *Chlamydomonas reinhardtii* was conducted as described previously with some modifications ^48^. The ClpP1-strep strain was inoculated into 24 L TAP liquid medium for 4 days under continuous light (40 μmol photo m^−2^ s^−1^). When cell numbers reached around 6×10^6^ cells/mL, the cells were collected by centrifugation at 3600 g for 6 min and resuspended in buffer A (20 mM Tris-HCl pH 8.0, 150 mM NaCl). The volume of the cell slurry was adjusted to 200 ml by the addition of 1 mM EDTA, 1 mM PMSF, and two tablets of EDTA-free protease inhibitor cocktail (Roche). Unless otherwise stated, subsequent steps were performed at 4◻. The cells were sonicated and centrifuged at 36,000 g for 30 min to remove debris. 1 mg/L avidin and 6 mM MgCl_2_ were added to the supernatant to improve the Strep tag binding efficiency with Strep-Tactin beads (Novagen). Next, the supernatant was further clarified by ultracentrifugation at 150,000 g for 1 h (Beckman Coulter rotor, 70Ti). Ammonium sulfate was slowly added to the supernatant at 25% saturation. After gently stirring for 30 min, the produced aggregates were removed by centrifugation at 36000 g for 20 min. Buffer E (20 mM Tris-HCl pH 8.0, 1 mM DTT, 10% glycerol) was added to the supernatant to dilute the ammonium sulfate concentration to 0.5 M and the solution was applied to pre-equilibrated hydrophobic column after passing through a 0.22 μm filter (Hitrap phenyl, GE Healthcare). Proteins were eluted by an ammonium sulfate gradient from 0.5 M to 0 M with ten column volumes. Clp-containing fractions were collected after visualization with SDS-PAGE and transferred to a Strep-Tactin gravity column which was pre-equilibrated with buffer B (20 mM Tris-HCl pH 8.0, 100 mM NaCl, 1 mM DTT). The protein was eluted with buffer C (buffer B+10%glycerol+2.5 mM Desthiobiotin (Novagen)) in four column volumes. The ClpP complex was concentrated and subjected to a Superdex-200 column pre-equilibrated with buffer D (20 mM Tris-HCl pH 8.0, 80 mM NaCl, 10% glycerol) and the desired fractions were collected. Purified ClpP complexes were concentrated to ~2 mg/ml by Amicon Ultra-15 Centrifugal Filter Units (Merck Millipore, Beijing China) with 100 kDa cut-off, supplemented with 10% glycerol and frozen at −80◻.

### Purification of recombinantly expressed proteins

#### EcClpP

The *EcClp*-pHUE plasmid was transformed into the *E. coli* BL21 (DE3) strain, and then transferred to 4 L lysogeny broth (LB) medium containing 100 μg/L ampicillin. When *E.coli* was grown to an OD 600 of ~ 0.6, 1 mM isopropyl β-D-1-thiogalactopyranoside (IPTG) was added to induce protein expression. After 4 hours, cells were collected by centrifugation at 4000 g. Unless otherwise stated, all purification steps were performed at 4◻. Cells were resuspended in lysis buffer (20 mM Tris-HCl, pH 8.0, 300 mM NaCl, 1 mM DTT, 10 mM imidazole, 1 mM phenylmethylsulfonyl fluoride (PMSF)) and lysed by sonication. Debris was removed by centrifugation at 36000 g for 40 min and the supernatant was passed through a 0.22 μm filter followed by transferring into the Ni-NTA gravity column. Protein was eluted with elution buffer (20 mM Tris-HCl, pH 8.0, 300 mM NaCl, 1 mM DTT, 250 mM imidazole) after two washes with buffer (20 mM Tris-HCl, pH 8.0, 300 mM NaCl, 1 mM DTT, 25 mM imidazole). Proteins in the eluted fractions were collected and digested with deubiquitinating enzyme USP2cc (1:50 molar ratio) to remove the ubiquitin-6×His tag. The digested solution was applied to Ni-NTA column again and the flow through was collected which contained EcClpP protein without tag. The collected protein was concentrated using Amicon Ultra-15 Centrifugal Filter Units (Merck Millipore, Beijing China) with 100 kDa cut off and frozen at −80 °C.

#### GroES, Cpn20 and Cpn11/20/23

All co-chaperonins were purified using the same method. Individual expression plasmid GroES-pET11a, Cpn20-pQlinkT or Cpn11/20/23-pQlinkT was transformed into *E. coli* strain BL21 (DE3). Transformants were picked and transferred to 6 L of LB medium containing 100 μg/L ampicillin. When cells were grown to an OD 600 of ~ 0.6, 1 mM IPTG was added to induce protein expression at 37◻ for 4 hours. Cells were harvested by centrifugation at 3600 g for 6 min. Unless otherwise stated, all subsequent steps were performed at 4◻. Cell pellets were resuspended in lysis buffer (30 mM Tris-HCl, pH 8.0, 60 mM NaCl, 1 mM DTT, 1 mM EDTA and 1 mM PMSF) sonicated. Lysates were centrifuged at 36,000 g for 30 min to remove cell debris. The supernatant was passed through a 0.22 μm filter followed by transfer to a pre-equilibrated source 30Q column (GE Healthcare) with 1 ml/min flow rate. Co-chaperonins were eluted by a linear salt gradient from 30 mM to 1 M NaCl with 10-fold column volume. Fractions containing protein according to UV absorption were separated by 15% SDS-PAGE and visualized by Coomassie staining. Eluted proteins were concentrated, then injected into a pre-equilibrated Superdex75 column (GE Healthcare) with a flow rate of 0.8 ml/min. According to UV absorption, the protein fraction was further analyzed with 15% SDS-PAGE and Coomassie staining. The pure co-chaperonin proteins were concentrated with Amicon Ultra-15 Centrifugal filters with 30 kDa cutoff, then flash-frozen in liquid nitrogen and stored at −80◻.

### Identification of CrClpP subunits by mass spectrometry

The protein subunits of the CrClpP complex were separated on an SDS-PAGE gel and visualized by Coomassie staining. Individual bands were cut out and digested by trypsin. Liquid chromatography-mass spectrometry (LC-MS) was done on a Thermo Scientific Q Exactive (QE) mass spectrometer at the Beijing Huada Protein R&D Center Co., Ltd. (Beijing, P.R.China). The Q Exactive mass spectrometry data were searched against the Phytozome v12.1(*Chlamydomonas reinhardtii*) database and NCBI-*Chlamydomonas* (taxid: 3052) database using 15 ppm peptide mass tolerance and 20 m/z fragment mass tolerance.

### Immunoprecipitation assays

Protein A-sepharose beads coupled with CrCpn20 or strep-tag antibodies were pre-equilibrated in lysis buffer containing 20 mM Hepes-KOH (pH7.5), 150 mM NaCl, 10 mM MgCl_2_, 20 mM KCl and 2 mM EDTA, then incubated with total protein or stroma protein which was prepared with the same lysis buffer under gentle stirring at 4◻. Protein A beads were washed three times with lysis buffer containing 0.1% Tween 20. Bound protein complexes were eluted with 2% SDS for 1 h at 4◻. The eluted proteins were separated by 12% SDS-PAGE and analyzed by immunoblotting.

### Analytical gel filtration

The ClpP interaction with co-chaperonins was analyzed by analytical gel filtration as described previously with some modifications ^38^. 1 μM ClpP and 2 μM co-chaperonin were incubated for 30 min at 4◻ in 20 mM MOPS-KOH, pH 7.5, 80 mM NaCl, 10 mM MgCl_2_, 10 mM KCl, 1 mM DTT, 10% glycerol. Then the protein complexes were loaded onto a Superdex 200 PC 3.2/10 column (GE Healthcare) at a flow rate of 0.05 ml/min. 50 μL fractions were collected and analyzed by immunoblotting with Strep-tag and Cpn20 antibodies.

### Asymmetric flow field-flow fractionation with multi-angle light scattering (AFFFF-MALS)

50 μg protein of purified CrClpP were loaded into a AFFFF-MALS device with a flow rate of 0.8 ml/min and a cross-flow rate of 2 ml/min using a 350 mm spacer and 10 kDa RC membrane (Wyatt Technology, Santa Barbara, CA, USA). The monitor methods employed a multiple-angle light scattering detector (DAWN HELEOS II, 658 nm; Wyatt Technology), a UV detector (1100 series, 280 nm; Agilent Technologies In., Santa Clara, CA, USA) and a differential refractive index detector (Optilab rEX, 658 nm; Wyatt Technology) ^49^. The CrClpP molecular weight was calculated in the presence of Dn/dc values equal to 0.185 ml/g.

### β-casein degradation assays

Degradation of the substrate protein β-casein was visualized by Coomassie Blue R staining after separating proteins via SDS-PAGE (12% acrylamide). The reaction mixture contained 20 mM Tris-HCl pH 8.0, 120 mM NaCl, 10 mM KCl, 10 mM MgCl_2_, 1 mM DTT, 10% glycerol, 0.4 μM EcClpP or CrClpP, 0.4 μM β-casein protein with or without 0.4 μM co-chaperonin (GroES, Cpn20 and Cpn11/20/23). 4 to 18 μM protease activator ADEP (#sc-397312, Santa Cruz) dissolved in DMSO was added to the reaction as indicated. The reaction was performed at 30°C, and the aliquots were taken at the indicated time points. The reaction was stopped by heating to 98°C for 10 min. Each reaction was performed at least three times and densitometric quantification of β-casein from the reactions was made with Photoshop CS6.

### Cryo-EM sample preparation and data collection

Holey carbon grids (Quantifoil R2/1, 200 mesh) were plasma cleaned using a Solarus plasma cleaner (Gatan), and an aliquote of 2 μl CrClpP sample was placed onto the glow-discharged grid. Then the grid was flash-frozen in liquid ethane by a Vitrobot Mark IV (Thermo Fisher Scientific). Movies were taken on a Titan Krios transmission electron microscope (Thermo Fisher Scientific) equipped with a Cs corrector and operated at an accelerating voltage of 300 kV with a nominal magnification of 18,000x (Table S3). Movies were collected by using a K2 Summit direct electron detector (Gatan) in super-resolution mode (yielding a pixel size of 1.318 A◻ after 2 times binning). Each movie was dose-fractioned into 38 frames and the exposure time was 7.6 s with 0.2 s for each frame, producing a total dose of ~38 e^-^ /Å^2^. The defocus value of the data set varied from −0.8 to −2.5 μm. We employed the SerialEM automated data collection software package to collect the images ^50^.

### Image processing and 3D reconstruction

A total of 8,064 movies were applied for CrClpP structure determination. Unless otherwise specified, single-particle analysis was mainly executed in RELION 3.1^51, 52^. All images were aligned and summed using MotionCorr2 ^53^ and CTF parameters were determined using CTFFIND4 ^51, 52^. We obtained 1,351,977 particles by automatic particle picking followed by manual checking, and 578,978 particles remained for further processing after reference-free 2D classification. Through one round of 3D classification, a ClpP-S1 dataset of 306,743 particles and a CrClpP-S2 dataset of 134,904 particles were obtained. Then multiple rounds of reference free 2D and 3D classification were applied to clean up the particles for each dataset. Classes with better structural features were combined and yielded ClpP-S1 dataset consisting 131,245 particles. After further Bayesian polishing and CTF refinement, a map at 3.3 Å resolution was obtained. Two classes with better structural features were combined and yielded ClpP-S2 dataset consisting 49,759 particles. After Bayesian polishing and CTF refinement, a ClpP-S2 map at 3.6 Å resolution was obtained. To improve the local resolution of the Cpn cap in the ClpP2-S2 map, the particles were subtracted by a soft mask focusing on the cap region and re-centered. We then applied 3D classification and obtained a cleaned-up dataset of 13,040 particles with better structural features especially more complete density of the Cpn mobile loops, which was further refined to 4.8 Å resolution of the Cpn11/20/23 map. ClpP-S2 map was then combined with the Cpn11/20/23 map together by using the *vopmaximum* function in Chimera ^54^, generating the composite ClpP-S2 map (termed ClpP-S2c). The overall resolution was determined based on the gold-standard criterion using an FSC of 0.143. The local resolution estimation was determined by Local resolution function in RELION 3.1.

### Model building of Clp subunits with the co-chaperonin

The structures of conserved regions of Clp subunits were predicted using the Rosetta server ^55, 56^. The predicted structures of conserved domains were docked rigidly into the density map in UCSF Chimera. The coordinates were further refined by the Real Space Refine module of the Phenix suite ^57^. On this basis of refinement, the Rosetta enumerative sampling method was applied to build the remaining residues of each Clp subunit *de novo* ^56, 58, 59^. The resulting model was adjusted manually in Coot ^60^. About the co-chaperonin Cpn11/20/23 model, each Cpn11, Cpn20, Cpn23 homology model was built with the tFold server (Tencent AI Lab) or Rosetta server . These models were docked into the cryo-EM map of ClpP-S2c using the fit in map command in UCSF Chimera ^54^. The resulting model was subjected to Rosetta and Phenix refinement ^57, 56^. The geometries and atomic model refinement statistics were evaluated by Molprobity in Phenix ^61^.

Cryo-EM data acquisition, 3D process information and model refinement statistics are summarized in Table S3. Figures were generated with either UCSF Chimera and ChimeraX ^54, 62^.

### Accession codes

Electron density maps have been deposited in the Electron Microscopy Data Bank under accession codes EMD-31171 for CrClpP-S1, EMD-31175 for ClpP-S2, EMD-31173 for ClpP-S2c and EMD-31174 for Cpn11/20/23. Related atom coordinates file also has been submitted to the Protein Data Bank, with accession codes 7EKO for CrClpP-S1, and 7EKQ for CrClpP-S2c.

## Acknowledgments

We are grateful to the staff of the NCPSS EM facility, Mass Spectrometry facility, and Database and Computing facility for instrument support and technical assistance. This work was funded by the Strategic Priority Research Program of Chinese Academy of Sciences (Grant No. XDA24020103-2, XDB37040103), the National Key Research and Development Program of China (2016YFD0100405, 2017YFA0503503) and the Ministry of Agriculture of China (2016ZX08009-003-005), the ‘Initiative d'Excellence’ program from the French State (Grant ‘DYNAMO’, ANR-11-LABX-0011-01), and the DFG (TRR 175, project C02). We thank Prof. Jean David Rochaix and Dr. Silvia Ramundo for their fruitful discussion.

## Author Contribution

C. L. and Y.C. supervised the project. N. W. executed all biochemical experiments. Y. W. and X. Z. collected the cryo-EM data. Y. W. did data processing with initial map from X. Z. Y.W. and N.W. did model building and structural analysis. Q. Z. started the project and optimized the protein purification. C. P. performed the MS analysis. W. Z. and Y. L. helped to purify protein. O. V. and M. S. were involved in the project design, data analysis and interpretation. C. L., N. W., O. V. and M. S. wrote the manuscript with modification from Y. W. and Y. C.

## Information

The manuscript contains five figures. The supplementary data including eight Figures and three Tables can be found enclosed with this article.

