## Supplemental tables and figures for "The cryo-EM structure of the chloroplast ClpP complex reveals an interaction with the co-chaperonin complex that inhibits ClpP proteolytic activity"

**SUPPLEMENTAL TABLE S1**

**Table S1. Summary information on peptides identified by mass spectrometry**

| Protein | Coverage (%) | Peptides | emPAI | Score | Mass (kDa) |
|---|---|---|---|---|---|
| ClpP1$_{H}$ | 49 | 182 | 5.53 | 6100 | 59.7 |
| ClpP1$_{N}$ | 32 | 20 | 1.12 | 608 | 19.9 |
| ClpP1$_{C}$ | 52 | 80 | 5.89 | 2334 | 23.1 |
| ClpP1$_{C}$’ | 52 | 67 | 4.56 | 1752 | 21.3 |
| ClpP4 | 73 | 655 | 1504.43 | 29289 | 38.7 |
| ClpP5 | 83 | 572 | 1100.42 | 27488 | 27.8 |
| ClpR1 | 51 | 437 | 19.56 | 17090 | 46.4 |
| ClpR2 | 62 | 370 | 786.48 | 16712 | 31.7 |
| ClpR3 | 68 | 709 | 725.04 | 26485 | 47 |
| ClpR4 | 77 | 416 | 266.30 | 15387 | 31.1 |
| ClpR6 | 62 | 487 | 384.00 | 21765 | 31.3 |
| ClpT3 | 34 | 8 | 1.23 | 381 | 31.5 |
| ClpT4 | 69 | 313 | 249.33 | 11991 | 26.8 |
| Cpn11 | 60 | 59 | 28.17 | 1103 | 13.5 |
| Cpn20 | 75 | 346 | 179.32 | 12531 | 22.8 |
| Cpn23 | 76 | 152 | 184.00 | 6229 | 25.8 |

**SUPPLEMENTAL TABLE S2**

**Table S2. Summary information on sequences solved in the structure. In column 4, the numbering is based on the mature protein, except for ClpP1$_{C}$ (based on ClpP1$_{H}$ precursor). Sequences solved up to the N- or C-termini are in bold letters.**

| Protein | N-terminal amino acids of the mature proteins | Length of precursor | length of mature proteins | Sequence present in structure |
|---|---|---|---|---|
| ClpP1$_{C}$[b] | $_{316}$NYLD | 524 | - | Ser345-Ala509 |
| ClpP4[b] | $_{50}$NSQP | 345 | 296 | Pro17-Asn195 |
| ClpP5[a] | $_{19}$ARRS | 256 | 238 | Pro42-Val234 |
| ClpR1[b] | $_{166}$AYGD | 411 | 246 | Tyr8-Lys224 |
| ClpR2[a] | $_{33}$KKSP | 282 | 250 | Cys41-Pro227 |
| ClpR3[a] | $_{33}$LVVH | 415 | 383 | Asp11-Ala348 |
| ClpR4[b] | $_{43}$RKLS | 293 | 251 | **Arg1**-Pro233 |
| ClpR6[a] | $_{46}$RRNV | 283 | 238 | Pro17-**Asn238** |
| ClpT4[b] | $_{65}$ATAT | 244 | 180 | **Ala1**-Pro162 |
| Cpn11[c] | $_{26}$AASV | 128 | 103 | Ala9-**Glu103** |
| Cpn20[c] | $_{23}$ATPV | 216 | 194 | **Ala1-Ser216** |
| Cpn23[c] | $_{27}$EAIS | 238 | 212 | **Glu1-**Gly139<br>Lys147-**Ala212** |

[a] Position of the transit peptide cleavage site predicted by PredAlgo

[b] Position of the transit peptide cleavage site determined by Derrien et al. (2012).

[c] Position of the transit peptide cleavage site according to Schroda (2014).

#### SUPPLEMENTAL TABLE S3

**Table S3. Cryo-EM data collection and refinement statistics.**

| | | ClpP | |
|---|---|---|---|
| **Data collection** | | | |
| EM equipment | | Titan Krios | |
| Voltage (kV) | | 300 | |
| Detector | | K2 Summit | |
| Pixel size (Å) | | 1.318 | |
| Electron dose ($e^-$/Å$^2$) | | 38 | |
| Exposure time (s) | | 7.6 | |
| Frames | | 38 | |
| Defocus range (μm) | | -0.8 to -2.5 | |
| **Reconstruction** | | | |
| Softwares | | Relion 3.1 | |
| Maps | ClpP-S1 | ClpP-S2 | Cpn11/20/23 |
| Final particles | 131,245 | 49,759 | 13,040 |
| Symmetry | C1 | C1 | C1 |
| Final overall resolution (Å) | 3.3 Å | 3.6 Å | 4.8 Å |
| **Atomic modeling** | | | |
| Softwares | Rosetta & Phenix & Coot | | |
| Structures | ClpP-S1 | ClpP-S2c | |
| Rms deviations | | | |
| Bond length (Å) | 0.009 (0) | 0.004 (0) | |
| Bond Angle (°) | 0.993 (21) | 1.049 (9) | |
| Ramachandran plot (%) | | | |
| Favored | 93.85 | 96.64 | |
| Allowed | 6.08 | 3.36 | |
| Outliers | 0.07 | 0 | |

#### SUPPLEMENTAL FIGURE LEGENDS

**Figure S1. Analysis of the CrClpP complex and its interaction with co-chaperonins**

Mass determination of purified CrClpP complexes by AFFFF. The horizontal blue line across the peak displays the molar mass.

**Figure S2. Analysis of the proteolytic activities of ClpP complexes from *E. coli* and *Chlamydomonas***

**(A)** Degradation of β-casein was monitored in reactions containing β-casein (16 μM), CrClpP (0.4 μM), EcClpP (0.4 μM), Cpn20 (0.4 μM), Cpn11/20/23 (0.4 μM) and ADEP dissolved in DMSO (4, 8 or 18 μM) as indicated. The reactions were performed at 30°C and aliquots taken at the indicated time points were analyzed via SDS-PAGE (15% gels) and Coomassie staining. The position of β-casein and the EcClpP protein are indicated. The casein degradation products resulting from proteolytic attack by the CrClpP complex are shown in the dotted box.

**(B)** Densitometric quantification of β-casein from the reactions with CrClpP and ClpP+GroES.

**(C)** Gel filtration of ClpP complexes in the presence or absence of ADEP. 1 μM ClpP, 2 μM co-chaperonin or 1 μM ClpP supplemented with 18 μM ADEP were injected into a Superdex 200 PC 3.2/10 column with 20 mM MOPS-KOH, pH 7.5, 80 mM NaCl, 10 mM $MgCl_2$, 10 mM KCl, 1 mM DTT, 10% glycerol. The relevant factions were collected and analyzed by western blot with Cpn20 antibodies.

**Figure S3. Data collection and processing of ClpP particles visualized by cryo-EM**

**(A)** Representative micrograph of CrClpP complexes. The scale bar equals 100 nm.

**(B)** 2D class averages of CrClpP complexes.

**(C)** Workflow of the 3D reconstruction based on cryo-EM data. A total of 578,978 particles were used for 3D classification.

**Figure S4. Statistics of the final density map of ClpP**

**(A)** Overview of ClpP-S2 particles.

**(B)** Overview of Cpn11/20/23 particles

**(C)** Gold standard Fourier Shell Correlation (FSC) curves of the final refinement map of ClpP-S1, ClpP-S2 and Cpn11/20/23.

**(D)** Local resolution map of ClpP-S1, ClpP-S2 and co-chaperonin densities.

**(E)** The particle orientation distributions in the final iteration of structure refinement. Red parts in the cylinders represent more particles in this direction.

**Figure S5. Alignment of the amino acid sequences of ClpP/R subunits**

Sequence alignment of ClpP and ClpR subunits from *Escherichia coli* (Ec), *Synechocystis* sp. PCC6803 (Sy), *Mycobacterium tuberculosis* (Mb), *Chlamydomonas reinhardtii* (Cr) and *Arabidopsis thaliana* (At), extracted form an alignment of 243 algal, plant and bacterial sequences. Structural elements derived from the crystal structure of *E. coli* ClpP are shown on top of the alignment (α: α-helices, β: β-strands). The three catalytic residues are marked by red arrows. The best-conserved residues are shown with a colored background. Chlamydomonas subunits (names in red) are shown with the experimentally-determined mature N-terminus boxed and sections not assigned in the Clp-S1 structure in faded colors. ClpP1_Cr is shown after removal of the insertion sequence IS1 (at pos 59, purple arrow), so the N-terminus of ClpP1c, starting near the end of IS1, is not shown. The less conserved regions of Clp subunits used for subunit assignment in Fig. S6B are underlined with red lines. The proline motif region is underlined with a blue line.

**Figure S6. Pseudo-atomic models of Clp subunits and close-up views of the fitting of long side chains of selected amino acids into the density map**

**(A)** Pseudo-atomic models of conserved sequences of Clp subunits. The sequences correspond roughly to amino acids 27 to 175 in EcClpP. Each color ribbon represents a different Clp subunit.

**(B)** Close-up views of the density maps accommodating long side chains of selected amino acids in individual ClpP/R subunits. The selected sequence regions are indicated in Fig. S5 and Fig. S6.

**Figure S7. Model-to-map fitting.**

**(A)** Density maps of Clp subunits. Ribbon presentations of the structural models of the individual subunits are docked into the cryo-EM densities with the subunit assignment indicated below. For each subunit, the N- and C-terminal residues of the assigned sequence are indicated.

**(B)** Correlation coefficient (CC) value of each subunit in the ClpP core complex. Individual Clp subunit was labeled in X-axis. The CC value is generated by Phenix 1.19.

**(C)** A superimposition of the subunit models (colored) and the cryo-EM density map (gray). Unassigned, additional density maps A2 and A3 are encircled with broken black lines.

**Figure S8. Properties of the ClpP-Cpn11/20/23 complex.**

**(A)** A superimposition of the EcClpP complex with ClpP-S2 particles. The additional density map A4 is the co-chaperonin complex.

**(B)** Correlation coefficient (CC) values of each subunit of the ClpP-Cpn11/20/23 complex. Individual Clp and co-chaperonin subunit was labeled in X-axis. The CC value is generated by Phenix 1.19.

**(C)** Density maps of co-chaperonin subunits. Ribbon presentations of the structural models of the individual co-chaperonin subunits, which are docked into the cryo-EM densities.

**(D)** Insertion of the mobile loops of Cpn20N into the hydrophobic clefts in the surface of the ClpP P-ring.

**Figure S1  Wang  et al**

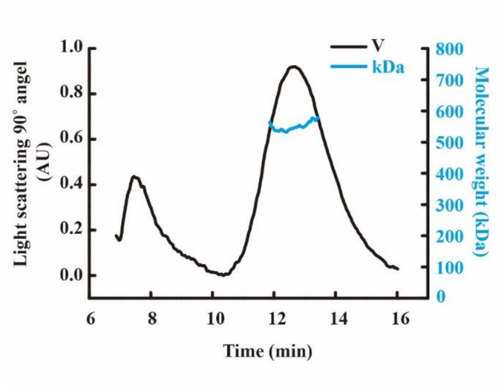

**Figure S2 Wang et al**

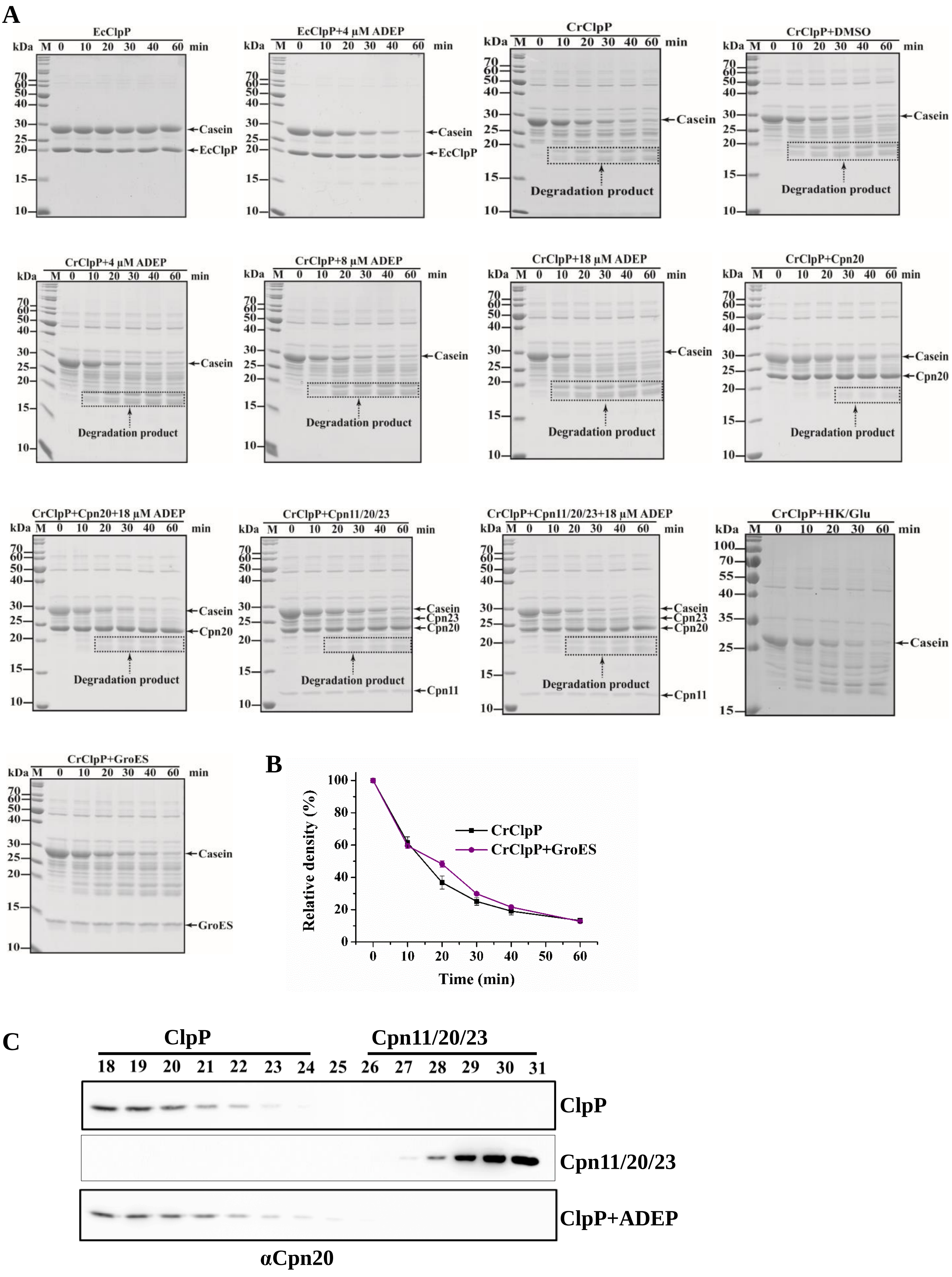

**Figure S3 Wang et al**

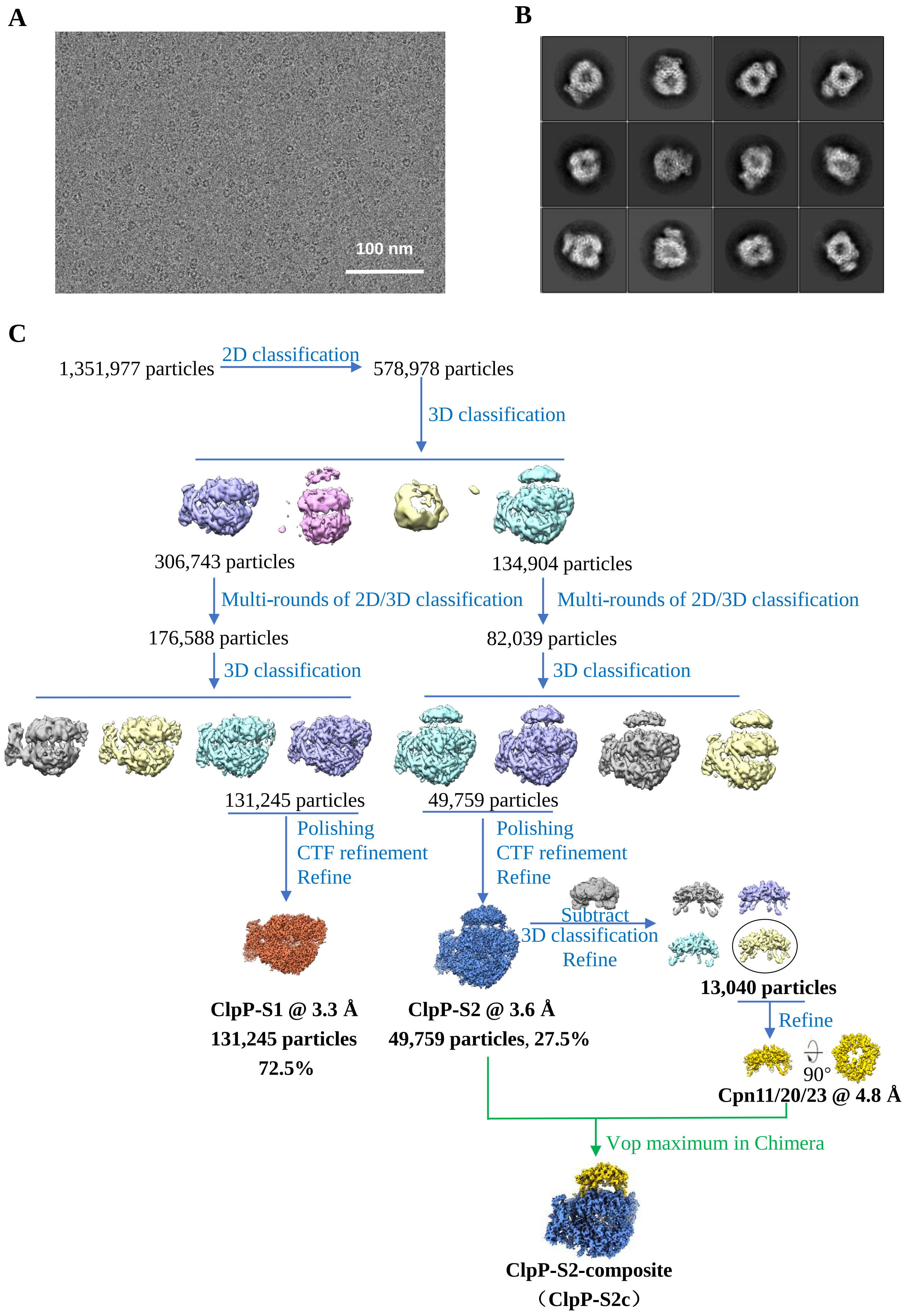

**Figure S4 Wang et al**

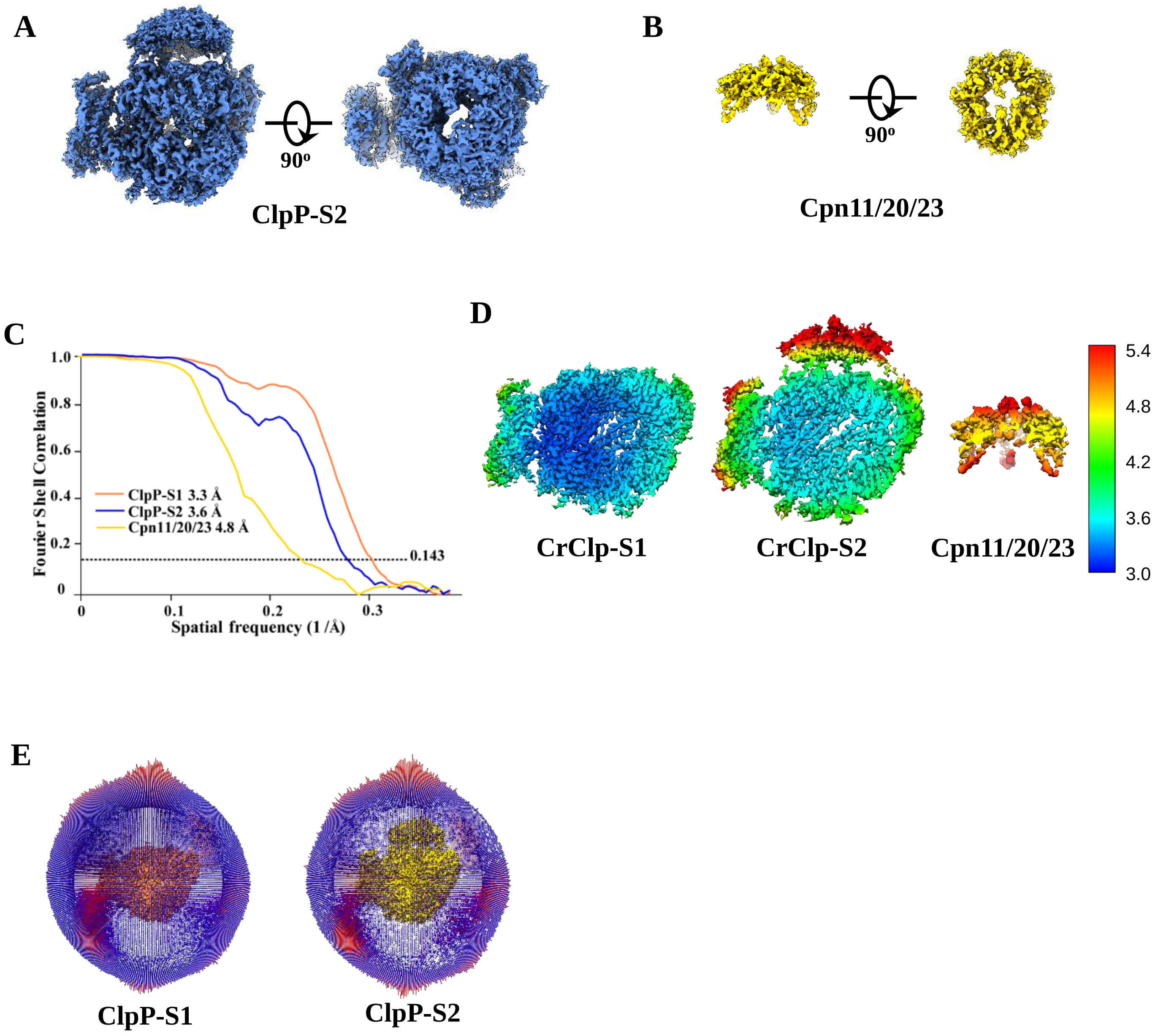

### Figure S5 Wang et al

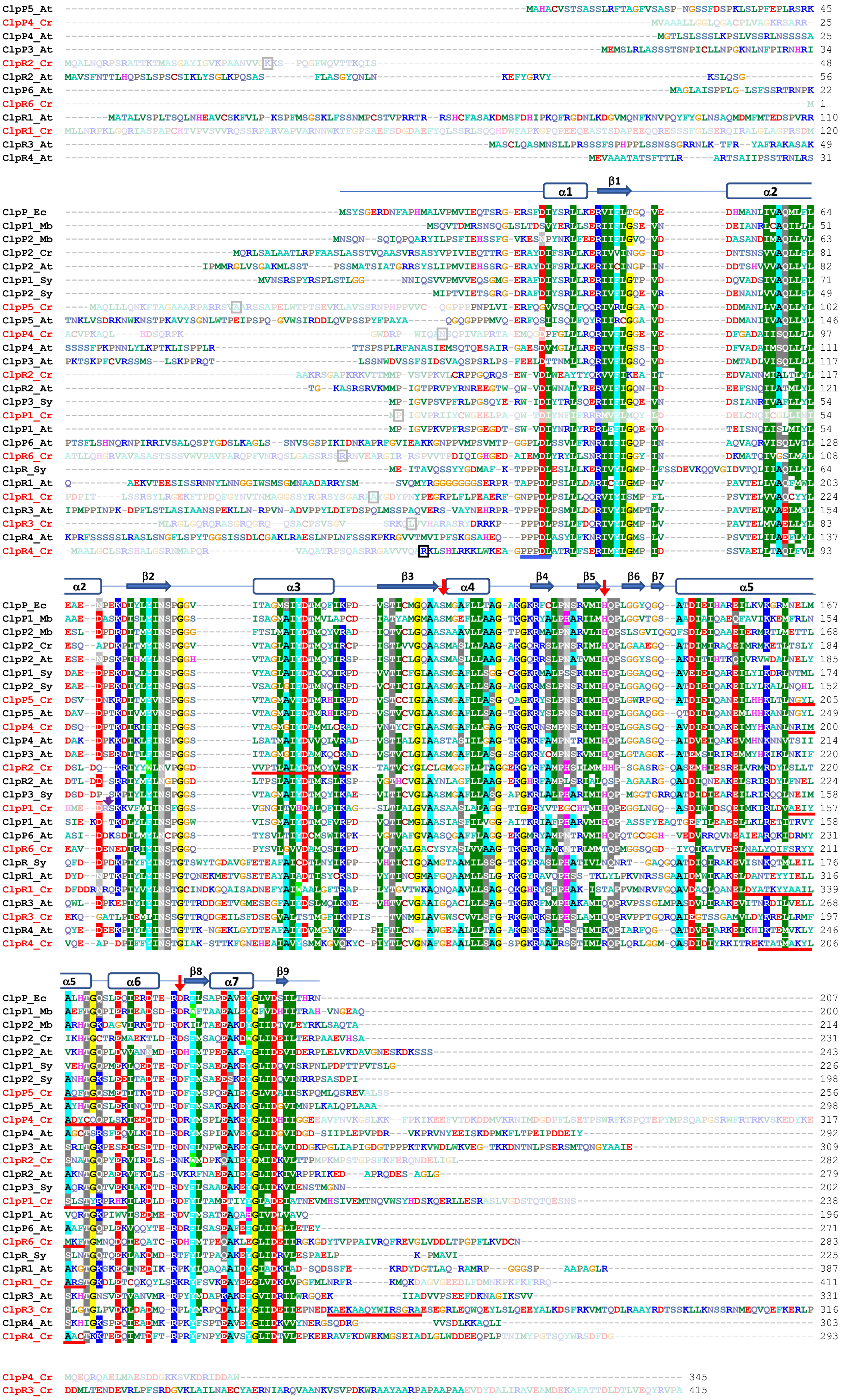

**Figure S6 Wang et al**

**A**

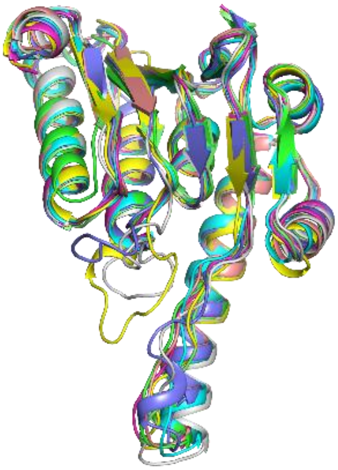

**conserved sequences**

**B**

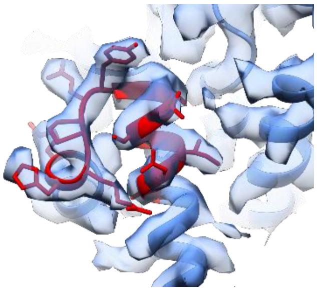

**$ClpP1_C$ AEIYSLSTYRPRHK**

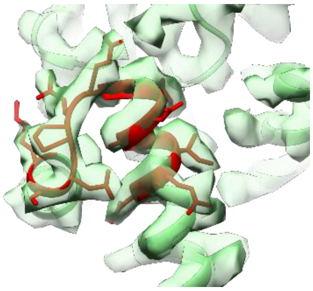

**ClpP4 NRIMADYCQQPLSK**

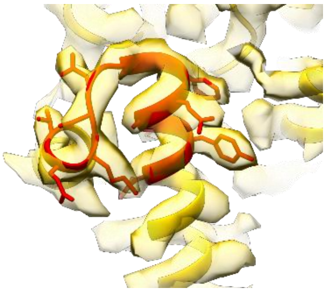

**ClpP5 NGYLAQFTGQSMET**

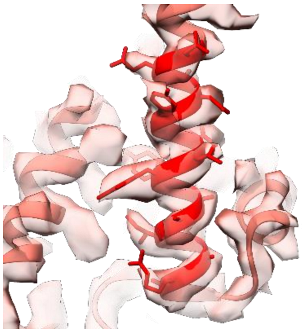

**ClpR1 DYATKYYAAILARS**

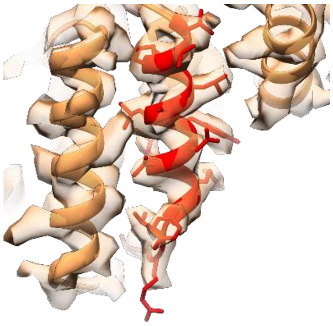

**ClpR2 VVPTLALYDTMQYVRS**

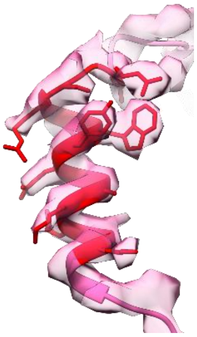

**ClpR3 KAEKAAQYWIRSGRA**

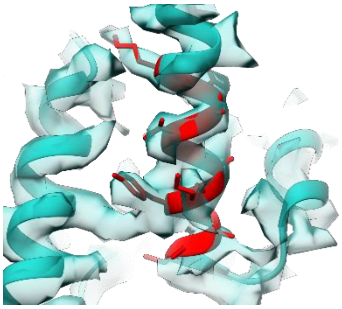

**ClpR4 KTATMAKYLAAC**

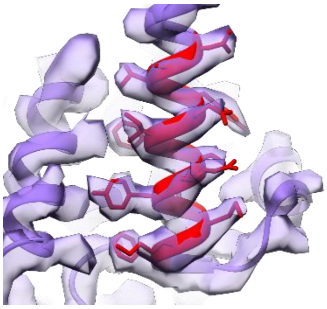

**ClpR6 NALYQIFSRYYMKF**

**Figure S7 Wang et al**

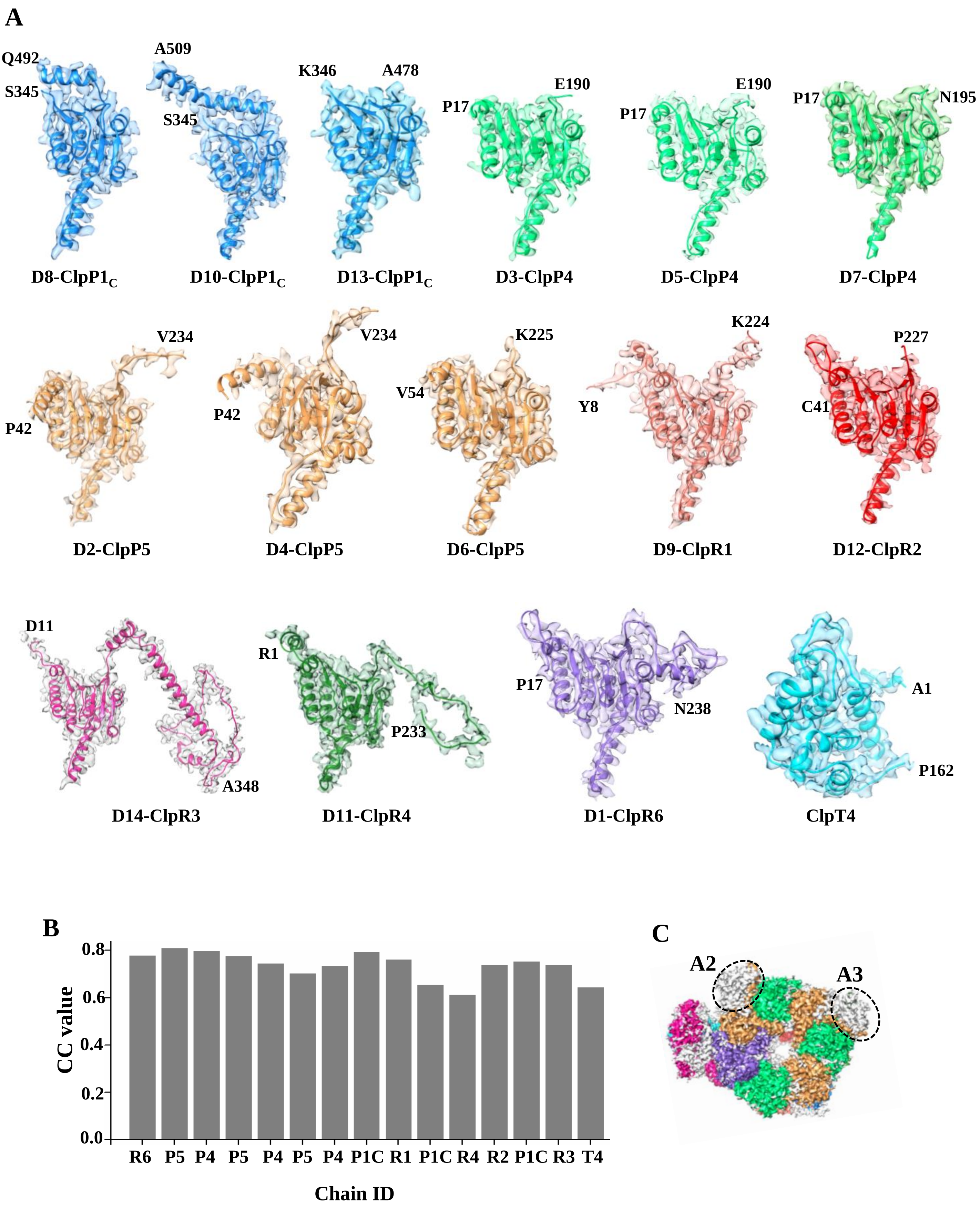

### Figure S8 Wang et al

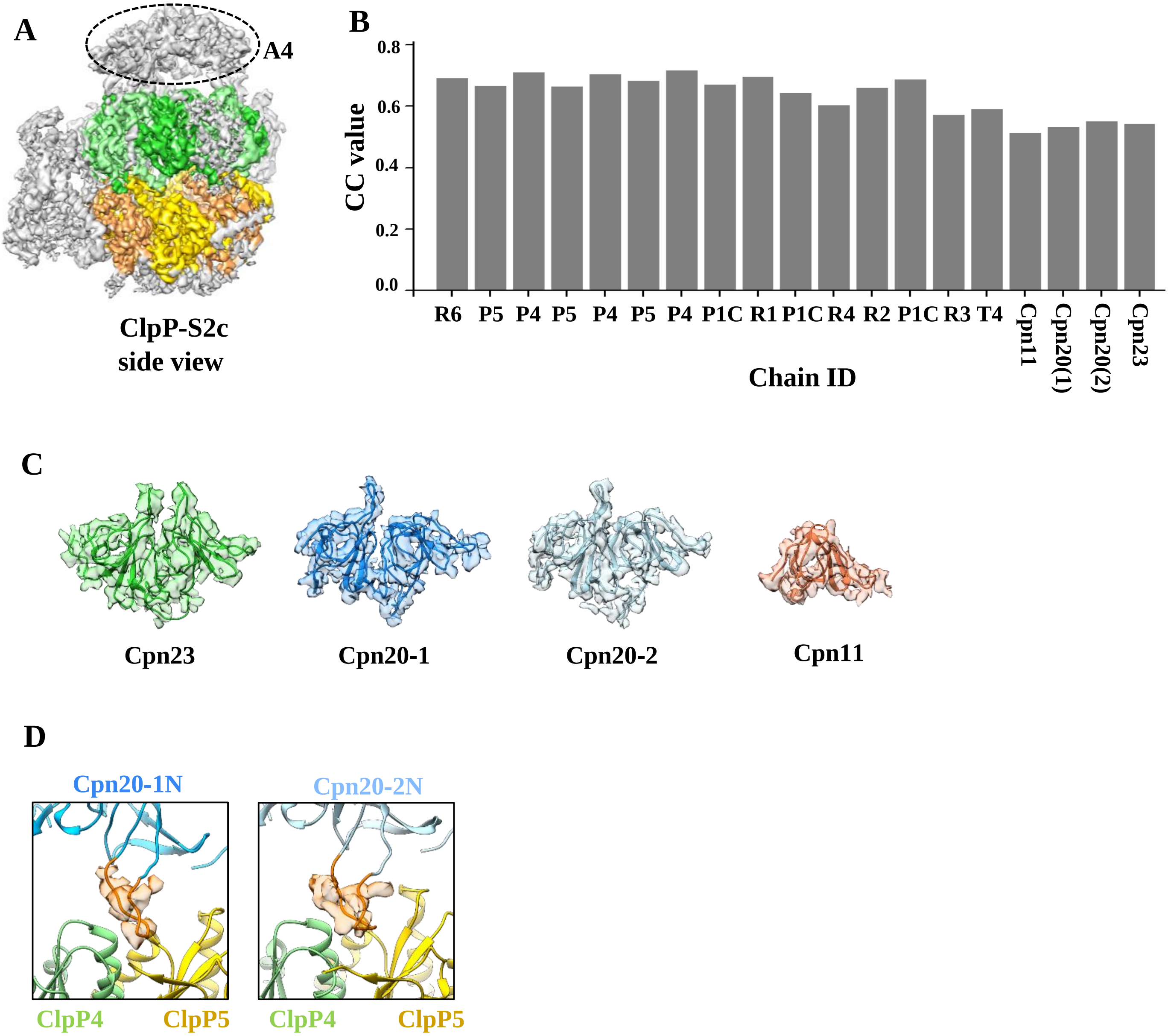
